# A Feature Learning Model Identifies Predictive Attributes of Mesenchymal Stromal Cell Efficacy

**DOI:** 10.1101/2025.10.20.683469

**Authors:** Sanique M. South, Yan Carlos Pacheco, Jay M. McKinney, Sara Bitarafan, Krishna A. Pucha, Nicholas M. Pancheri, Kaitlyn Link, Angela Lin, Levi B. Wood, Nick J. Willett

**Affiliations:** Phil and Penny Knight Campus for Accelerating Scientific Impact, Department of Bioengineering, University of Oregon; Eugene, Oregon, USA; Research Division, VA Medical Center; Atlanta, Georgia, USA; Department of Orthopaedics, Emory University; Atlanta, Georgia, USA; Wallace H. Coulter Department of Biomedical Engineering, Georgia Institute of Technology and Emory University; Atlanta, Georgia, USA; Department of Medicine, Division of Cardiology, Emory University; Atlanta, Georgia, USA; Parker H. Petit Institute for Bioengineering and Bioscience, Georgia Institute of Technology; Atlanta, Georgia, USA; George W. Woodruff School of Mechanical Engineering, Georgia Institute of Technology; Atlanta, Georgia, USA

## Abstract

The therapeutic efficacy of human mesenchymal stromal cells (hMSCs) is highly variable, limiting their clinical translation for musculoskeletal diseases and other regenerative medicine applications. There is a poor understanding of the critical quality attributes correlating to therapeutic efficacy of hMSCs. To address this challenge, we analyzed pre-clinical *in vitro* secretome profiles and *in vivo* therapeutic efficacy of hMSCs from multiple human donors. hMSCs from different donors showed significant differences between donors in therapeutic efficacy when assessed in a rat post-traumatic osteoarthritis (OA) model. A partial least squares feature learning model was trained to evaluate differences between more and less therapeutic donor hMSCs by examining cytokine secretion profiles, to predict donor-specific therapeutic outcomes. More therapeutic hMSCs exhibited increased secretion of GM-CSF, GRO, IL-4, and PDGF-AA, whereas less therapeutic donors had higher TNF-α, IL-6, and MCP-1 secretion. The cytokine profile was accompanied by evaluation of MAPK pathway, which revealed distinct differences in phospho-protein signaling between more and less therapeutic hMSC secretome profiles. Pharmacological inhibition of JNK signaling in more therapeutic donor cells decreased hMSC secretion of the key therapeutic associated cytokines and shifted hMSC secretome towards a less therapeutic profile. Prospective validation of cells from additional donors demonstrated significant correlations between predicted and observed pre-clinical in vivo efficacy to attenuate OA. This approach identifies critical quality attributes enabling consistent prediction of therapeutic potency, thereby addressing a major barrier to scalable and effective cell therapies. These findings advance precision cell-based therapies and offer a framework for standardized donor screening in clinical applications.

**Summary:** A feature learning model was developed, trained, and validated to identify critical quality attributes of MSCs that predict therapeutic potency.

## INTRODUCTION

Cell therapies have emerged as potentially transformative approaches in modern medicine, with applications spanning regenerative medicine, immunomodulation, and tissue repair. Among these therapies, mesenchymal stromal cell (MSC)–based therapies have garnered particular attention due to their multipotent differentiation capabilities and robust immunomodulatory functions. MSCs are under investigation for a wide array of conditions including cardiovascular diseases, autoimmune disorders, and musculoskeletal injuries, with joint injuries and osteoarthritis (OA) representing one of the most common indications (*1, 2*). Despite extensive preclinical data demonstrating regenerative benefits and anti-inflammatory effects of MSC-based therapies, clinical trials investigating use for OA and other indications have often yielded discordant results (*2–4*). A recent randomized clinical trial compared different MSC-based cell products and found that there was no overall benefit of cell therapy 1 year post-treatment compared to gold standard corticosteroid injections (*2*). In light of widespread clinical use of MSC-based therapies, the FDA has issued consumer alerts and public warnings on the use of unapproved stem cell therapies (*5, 6*). Scientific societies, like The American Society for Bone and Mineral research (ASBMR) (*7*) and International Society for Cell and Gene Therapy (ISCT), have published perspective articles stating the need for rigorous scientific evidence supporting clinical use of these cell therapies (*7*). There is a critical need in the cell therapy field for products with consistent potency, rigorous science supporting best practices for use, and strategies that can reliably predict therapeutic outcomes.

Achieving consistent potency in MSC therapies is essential for the successful translation into clinical practice. Therapeutic effects of MSC-based therapies have traditionally been thought to be mediated either through direct engraftment and tissue regeneration by exogenously delivered cells, or through paracrine mediated activity, which then induces a response from the host cells and tissues. Recent studies, including our own work, have provided evidence that the majority of exogenously delivered MSCs do not maintain viability long term; instead, their therapeutic benefits appear to be mediated predominantly through the secretion of paracrine factors (*8–10*). Consequently, differences in intracellular signaling and cytokine secretion profiles between MSC donors, such as divergent levels of growth factors and immunomodulatory molecules, are implicated in the overall therapeutic potency of a given patient or donor cells.

A major challenge in the field of cell therapy is the inherent heterogeneity of MSCs. Even when isolated using standard criteria, MSCs isolated from different donors exhibit significant variability in their secretome and functional attributes(*11, 12*). Pre-clinical studies typically utilize cells derived from a single donor which are subsequently expanded and utilized for multiple studies; yet, in clinical utilization, most current approaches utilize autologous cells where patient to patient variability in the cells could contribute to inconsistent therapeutic efficacy (*13, 14*). Pre-clinical *in vivo* studies investigating MSCs for use as OA therapeutics have not addressed this donor heterogeneity or variability. Furthermore, among the majority of clinical trials using MSC treatment for OA the standard approach involves the isolation, expansion, and autologous injection of hMSCs, with no screening metrics implemented to assess quality or potency of the cellular therapy (*11, 15*). Current potency assays such as IFNγ/TNFα stimulation, angiogenesis, and T-cell immunomodulation tests often rely on singular, binary readouts and are not readily integrated into manufacturing pipelines (*16, 17*). These methods, while useful in demonstrating variability among donor cells, lack the predictive power needed to ensure reliable and consistent clinical efficacy.

The immunomodulatory and anti-inflammatory paracrine signaling properties of hMSCs have been linked to their biological activity and have recently been considered an essential quality attribute of hMSC therapeutic potency for OA therapy (*18–20*). Based on this premise, the aim of this study was to identify secreted paracrine factors related to therapeutic efficacy of hMSC donors. To address this aim, we quantified heterogeneity in therapeutic activity among four hMSC donors using the rat medial meniscal transection (MMT) pre-clinical model of OA, measuring regenerative and therapeutic outcomes. We then quantified cytokines and growth factors secreted by these cells under culture conditions, identifying a targeted paracrine profile associated with enhanced therapeutic outcomes. Next, we used RNAseq and multiplexed phospho-protein signaling panels to gain insight into potential mechanisms responsible for differences in paracrine factor secretion. Using secreted cytokines as training data, we established a feature learning model relating our *in vitro* and *in vivo* data to identify critical quality attributes of hMSCs that predict their therapeutic potency. This model was built using partial least squares regression (PLSR) to prospectively predict therapeutic efficacy of hMSC donors based on their secretion of immunomodulatory cytokines during culture. The accuracy of the feature learning model predications was then evaluated in a prospective validation study testing the efficacy of the new hMSC donors in the pre-clinical OA model. The validation studies confirmed significant correlations between predicted and observed therapeutic efficacy. Collectively, these data establish a robust framework for hMSC donor screening of cellular quality attributes promoting consistent therapeutic outcomes and advancing successful implementation of cell therapies in clinical translation.

## RESULTS AND DISCUSSION

### A feature learning model framework of hMSC therapeutic potency

Cellular therapies have generally shown great promise in pre-clinical studies; however, clinical trials have shown highly variable results. This variability is thought to be in part from the high heterogeneity in cell potency between donors as well as variability in the host environment. Understanding the reliability and potency of donor hMSCs is a critical step to ensure consistent and optimized therapeutic outcomes for disease therapy. We developed a feature learning model framework integrating our *in vitro* and *in vivo* data to identify critical quality attributes of hMSCs that predict their therapeutic potency. The objective of this framework was to first generate a predictive model of hMSC therapeutic potency based on hMSC secretome data and therapeutic efficacy data from a pre-clinical post traumatic OA (PTOA) model (Model Development, Figure 1).

**Figure 1.**
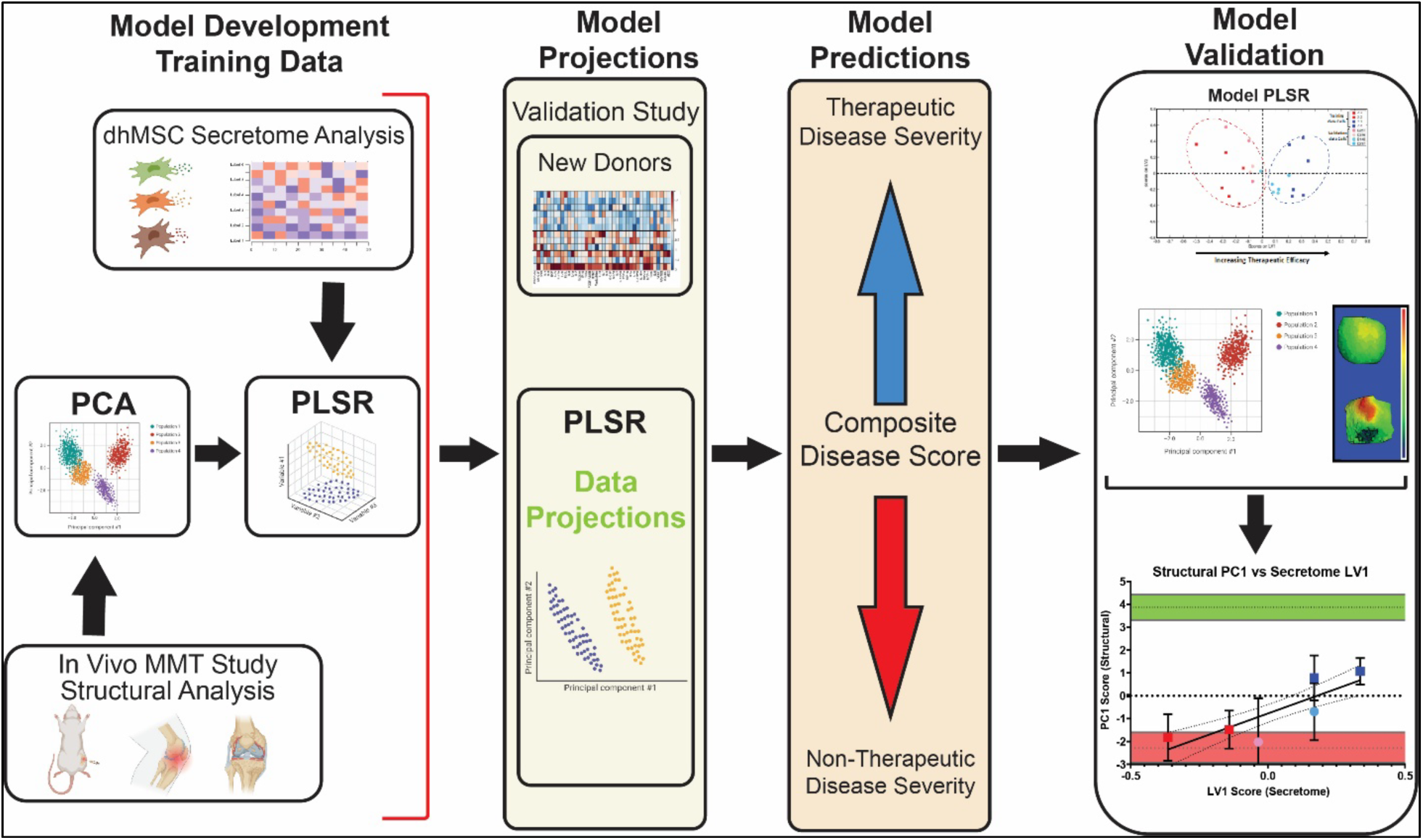
Overall preclinical experimental design of our feature learning model of hMSC therapeutic potency. The figure illustrates the four major components of the feature learning model and how the model was validated. The model includes model development, model projections, model predictions and model validation with additional studies referred to as the validation data (figure created in BioRender.com).

Next, we projected additional donor hMSC secretome profiles onto the PLSR model and determined the predicted composite disease scores indicative of therapeutic efficacy (Model Projection & Predictions, Figure 1). Less and more therapeutic donor hMSCs were identified and hMSCs from each group were validated in our preclinical PTOA model. In addition, we conducted a correlational analysis between donor hMSC secretome scores and structural outcome parameter scores (Model Validation, Figure 1). Overall, these data establish a robust framework to screen hMSCs based on exhibited critical cellular attributes to predict therapeutic potency and to promote consistent disease therapy outcome and advance cell therapy efficacy in clinical translation.

### hMSCs from different donors yield variable therapeutic outcomes in a rat OA model

Effective clinical translation of cell therapies has been limited by factors including high variability and heterogeneity of MSCs and poor understanding of critical quality and potency attributes (*15, 21*). To identify hMSC cellular attributes that relate to therapeutic outcomes donor heterogeneity was assessed *in vitro* and *in vivo* using four unique hMSC donors sourced from Emory Personalized Immunotherapy Core (EPIC) at Emory University and RoosterBio. All hMSCs used for this study were from young, healthy patients, isolated from their bone marrow, and expanded to passage 4 to minimize variability from factors other than donor-specific differences. A well established *in vivo* preclinical rat model (6-week medial meniscal transection (MMT)) was used to assess therapeutic potency. This is a clinically relevant model that mimics key clinical features of OA symptoms and disease progression. Cells from an individual donor or a saline control were injected intra-articularly 3 weeks post-MMT surgery, a time point associated with early OA development(*22*). Six weeks after surgery, tissues were collected, and a detailed quantitative analysis was performed of articular cartilage changes in established OA using various morphological parameters for the total and medial 1/3 of the medial tibial condyle.

Quantitative analysis of the articular cartilage, using microCT revealed distinct difference between the sham and other treatment/injury groups (Figure 2). The data showed that donors 3 and 4 yielded improved therapeutic outcomes, relative to hMSC donors 1 and 2, including significantly attenuated increases in articular cartilage volume and thickness, fibrillation development (surface roughness), and development of lesions (lesion volume) (Figure 2 A – D). Total and medial cartilage volume, mineralized osteophyte and lesion volume total were significantly lower in the donor 3 and donor 4 treated groups in comparison to the MMT-Saline group (Figure 2A). For total articular cartilage volume, donors 3- and 4- treated groups showed significantly reduced articular cartilage volume relative to the saline control group, while donor 1- and 2- treated groups showed no difference in volume compared with the saline group (Figure 2B). Similar outcomes were observed in the medial 1/3 region for articular cartilage volume (Figure 2E). For total articular cartilage surface roughness, donors 3- and 4- treated groups showed significantly attenuated articular cartilage surface roughness relative to MMT/Saline; however, donors 1 and 2 showed no difference in surface roughness with the MMT/Saline disease control (Figure 2C, 2F). Donor 3 and 4 treated groups had reduced total articular cartilage lesion volume in comparison to MMT/Saline group (Figure 2D). In the Medial 1/3, no differences were observed between MMT saline and the MMT/hMSC groups for exposed bone area (Figure 2G). While donors 3 and 4 treated groups showed a trend towards less exposed bone area relative to the MMT/Saline (Figure 2D, 2G) this was not statistically significant. Representative contrast enhanced microCT and histological images qualitatively show the lesions found in the MMT/Saline, donor 1 and donor 2 treated groups in comparison to the sham (Figure 2H - K); representative images from donor 3 and donor 4 treated groups showed inhibition of cartilage degradation with no lesions evident in the cartilage surface (Figure 2L, 2M). Similarly, additional structural outcomes of the full medial and 1/3 medial section of the tibia shows reduced osteophyte volume in donors 2, 3, and 4 treated groups relative to the MMT/Saline group (Figure S1). In addition, all MMT/hMSC groups displayed lower subchondral bone volume and thickness compared to the MMT/Saline group (Figure S1). Overall, these metrics revealed a spread in the therapeutic efficacy of MSC derived from different donors, with donors 3 and 4 yielding improved (more) therapeutic outcomes compared to poorer outcomes from donors 1 and 2 (less).

**Figure 2.**
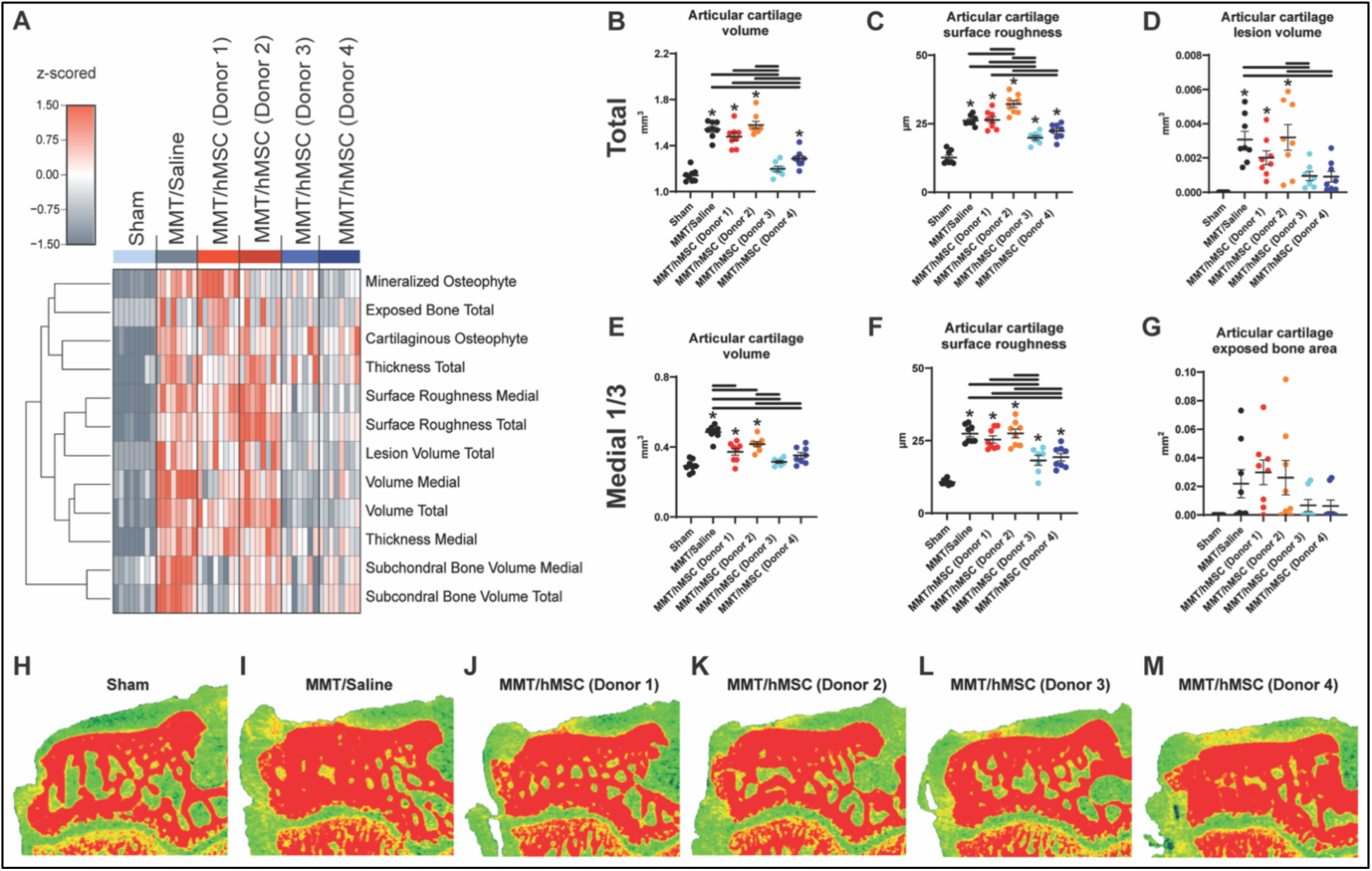
Effects of hMSCs on articular cartilage therapeutic structural outcomes in the total and medial 1/3 of the medial articular cartilage in MMT joints. (A) Represents the total and medial 1/3 region of the tibial plateau structural parameters analyzed by microCT. Quantitative analysis results from the total (B) articular cartilage volume, (C) articular cartilage surface roughness and (D) articular cartilage lesion volume. Similarly quantitative analysis was completed on the medial 1/3 region of the tibial plateau articular cartilage (E) articular cartilage volume, (F) articular cartilage surface roughness and (G) articular cartilage exposed bone area. (H-M) Representative contrast enhanced microCT images highlights the structural difference in MMT groups compared to sham. Lesions are seen in the MMT/Saline, MMT/hMSC donor 1 and MMT/hMSC donor 2 while MMT/hMSC donor 3 and MMT/hMSC donor 4 showed no lesions. Data presented as mean +/− SD. n=7 for MMT/hMSC donor 3 and n=8 for all other groups. * Represents significant differences (p < 0.05) between individual MMT groups relative to sham. ‘__’ Bars indicate significance (p < 0.05) between MMT groups.

### hMSC secretion of GM-CSF, GRO, and IL-4 relate to OA therapeutic outcomes

Inflammation has been well-characterized as playing a key role in OA pathogenesis (*23*). To recapitulate an *in vivo* inflammatory OA microenvironment *in vitro,* hMSCs were conditioned with IL-1β. IL-1β was used because it serves as a major pro-inflammatory cytokine in OA and many other clinical conditions (*24*). To assess differences in hMSC paracrine signaling between more therapeutic (donors 3 and 4) and less therapeutic donors (donors 1 and 2, Figure 1), we used a multiplexed immunoassay to quantify 41 cytokines and growth factors secreted into the medium by each donor during a 24 hour culture period (Figure 3A). The more therapeutic hMSCs (donors 3 and 4) secreted elevated levels of a few targeted factors, while the less therapeutic hMSCs (donors 1 and 2) secreted elevated levels of the majority of the cytokines and growth factors within the panel (Figure 3A). We accounted for the multi-dimensional nature of the data by using a partial least squares discriminant analysis (PLSDA) to identify a profile of cytokines that distinguished more therapeutic (donors 3-4) from less therapeutic (donors 1-2) donors (Figure 3B). The analysis identified a latent variable (LV1), which separated more therapeutic donors (3 and 4) to the right and less therapeutic donors (1 and 2) to the left. The LV1 consisted of a profile of cytokines associated with the more therapeutic (positive) or less therapeutic (negative) cells (Figure 3C). Univariate analysis revealed significant upregulation of granulocyte macrophage colony stimulating factor (GM-CSF), chemokine ligand 1 (GRO), and interleukin (IL)-4, in more therapeutic cells (donors 3 and 4) compared to less therapeutic (donors 1 and 2) (Figure 3D). Relating *in vitro* hMSC cytokine secretion with *in vivo* therapeutic outcomes demonstrated that hMSC donors which exhibited more therapeutic effects *in vivo* yielded increased secretion of key chemokines (GM-CSF and GRO), cytokines (IL-4) and growth factors (PDGF-AA) *in vitro*.

**Figure 3.**
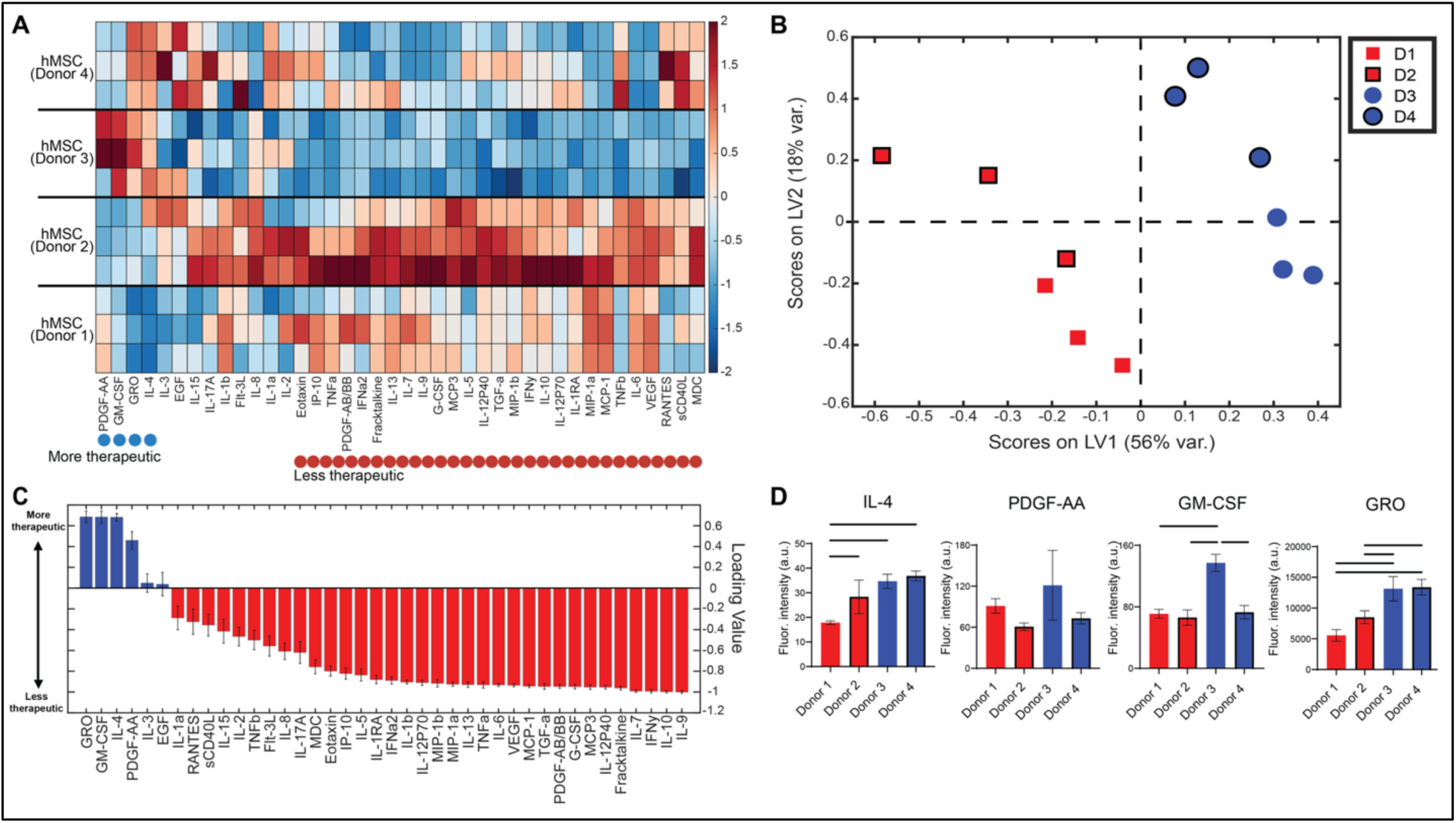
Paracrine signaling response of hMSCs in an IL-1β OA simulated microenvironment. Multiplexed immunoassay analysis of 41 cytokines (columns; z-scored) secreted from hMSCs in IL-1β conditioned media. (A) Donors 1 and donor 2 demonstrated overall increased cytokine secretion levels relative to donor 3 and donor 4. (B) PLSR analysis identified a profile of cytokines along LV1 that identified a distinct separation between less therapeutic (red; left side of figure) and more therapeutic hMSCs (blue; right side of figure). Variability accounted for in each LV is included on respective axes labels. (C) Loadings plot demonstrating relative contribution of cytokines to PLSR scores obtained show that the cytokines GM-CSF, GRO, IL-4, and PDGF-AA contribute to separating out more therapeutic hMSC donors while all other cytokines assessed contribute more to less therapeutic hMSCs. (D) IL-4, GM-CSF, and GRO were assessed independently and demonstrated significantly higher secretion levels while PDGF-AA levels were trending higher in more therapeutic hMSC donors (donor 3 and donor 4) relative to less therapeutic donors (donor 1 and donor 2).

The key MSC secreted cytokines and growth factors identified are known to have critical roles in joint health and OA development and progression. GM-CSF has been shown to enhance the mobilization of MSCs from the bone marrow and increased MSC secretion of GM-CSF has demonstrated potentiated therapeutic efficacy of articular cartilage repair induced by microfracture (*25, 26*). The chemokine GRO has been shown to initiate cartilage degradation and potentiate inflammation in joint tissues; these resulting cellular actions align with the well-defined role of GRO in trafficking monocytes and neutrophils to sites of injury (*27, 28*). Secretion of GRO would not necessarily result in increased degradation in the joint space as there are important distinctions to be drawn between the chronic nature of the OA inflammatory environment and an acute response induced by the hMSC secretome. To resolve chronic inflammation, an acute event or activation is often needed to recruit a new wave of immune cells and activate different inflammatory cascades to resolve inflammation and induce a pro-regenerative response (*29*). This mechanism is common in other chronic inflammatory and wound healing environments where acute inflammatory events are necessary to resolve chronic inflammation and transition to pro-regenerative immune responses to regulate inflammation (*30–32*). GRO induced trafficking of monocytes and neutrophils may be contributing to an acute event rather than potentiating the chronic inflammatory state of OA with more sustained inflammation.

In addition to secretion of chemotactic factors, the hMSCs from the more therapeutic donors demonstrated potentiated secretion of the potent immunomodulatory cytokine, IL-4 (*33*). IL-4 has demonstrated chondroprotective effects in pre-clinical models of OA, where MSC spheroids transduced with IL-4 yielded better cartilage protection and pain relief, relative to naïve MSCs *in vivo* (*34–37*). Further studies demonstrated that IL-4 can protect cartilage by inhibiting inducible nitric oxide synthase (iNOS) and NO which yields subsequent suppression of IL-1β and TNF-α (*38–41*).

The growth factor PDGF-AA has been studied extensively for its ability to yield therapeutic outcomes for OA treatment, both in combination and independently of MSC therapeutic delivery (*42*). PDGF has both tissue regeneration and anti-inflammatory properties and has been implicated as a critical component of platelet rich plasma (PRP) therapies (*42, 43*). In pre-clinical studies PDGF overexpressing MSCs were shown to exert anti-fibrotic, anti-inflammatory, and pro-chondrogenic capacities in a canine model of OA, through both the reduction of pro-inflammatory factors and MMP-13 (*44*). While the role of each of these specific cytokines and growth factors have been implicated in MSC therapeutic action in OA this profile of secreted factors is unique and identifies a potential signature or cocktail for development of a novel combinatorial therapeutic, accounting for all hMSC cytokines identified. This profile of secreted factors is specific to a more therapeutic profile and illustrates a target of secreted factors needed to achieve beneficial outcomes. Overall, these data show that there are significant differences in paracrine signaling between hMSCs with more or less *in vivo* therapeutic efficacy.

### Transcriptional profiling identifies immune and OA-related transcriptional signatures associated with therapeutic efficacy

Having identified a secreted cytokine profile associated with the therapeutic efficacy of hMSC donors, we next investigated the related transcriptional signatures from each hMSC donor after 24 hours of culture in the OA simulated microenvironment. Bulk RNA-seq was conducted to identify transcriptional signatures related to therapeutic efficacy. Using Hierarchical clustering of the 3,489 genes remaining after filtering (Methods), we identified distinct clusters of genes associated with the more and less therapeutic donor hMSC groups (Figure 4A). These data revealed that donors with different therapeutic efficacy (more vs. less therapeutic) have distinct gene expression profiles. Differential expression analysis identified 1,523 differentially expressed genes (DEGs) between the more and less therapeutic groups (722 genes up-regulated in more, 801 genes down-regulated in More) (Figure 4B, Sup Table 2). To gain insight into the functional implications of the identified DEGs, we applied gene set variation analysis (GSVA) to identify genes that were enriched in 2,198 canonical gene sets (Figure 4C) (*45*). To assess the relationship between transcriptional signatures and therapeutic outcomes, we used Pearson correlations to regress transcriptional pathways against therapeutic scores computed from the PCA analysis on microCT measurements (First principle component, PC1). This analysis identified signaling pathways that were both significantly correlated with therapeutic scores and different between More and Less therapeutic groups. These included MAPK, PTEN and Notch, which are key regulators of cytokine and growth factor expression (Figure 4, Sup Fig S2-S3, Sup Table 2). Additionally, gene sets related to pro-inflammatory cytokines were positively correlated with more therapeutic cells, whereas gene sets related to interleukins −12, −27, and −37 were all significantly correlated with less therapeutic cells. Less therapeutic cells were also correlated with OA-related genes, including collagen and integrin pathways. Together, these data define key transcriptional differences between more and less therapeutic donor cells associated with immune signaling and OA-related pathways.

**Figure 4.**
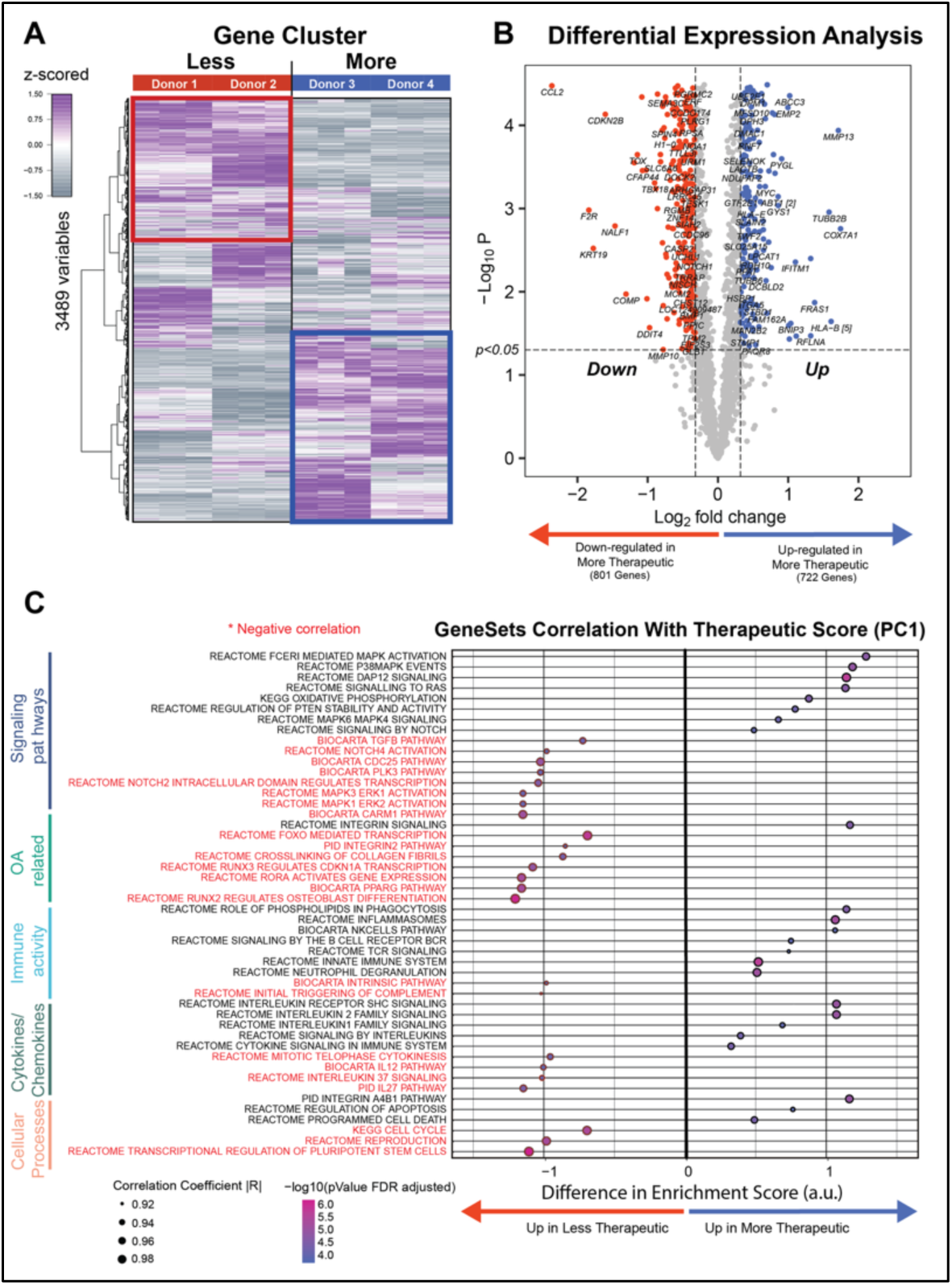
RNAseq analysis of hMSC cells reveals transcriptional signatures associated with therapeutic scores. (A)Transcriptional profile of 3,489 genes identified distinct patterns of gene expressions between less and more therapeutic groups (N=3/MSC donor, data are z-scored). (B) Comparison of more therapeutic gene expression profile to less therapeutic group, resulted in 1523 differentially expressed genes (DEGs) in hMSC where upregulated genes are shown in blue and down regulated genes are in red (|log_2_ fold change| ≥ 0.32, unadjusted p<0.05). (C) Pearson’s correlation coefficient of therapeutic score (PC1) versus gene sets enrichment score identified by GSVA (rows) in hMSC. The size of dots represents absolute value of the correlation coefficient (|R|) and the ones with negative correlation are noted with red asterisk (*). The color bar represents p-values (FDR adjusted) associated with the correlation slope.

### Inhibition of JNK activity reduced secretion of GM-CSF, GRO, IL-4, and PDGF-AA in more therapeutic hMSCs

The MAPK and JNK pathways have been shown in prior studies to be upstream regulators of many critical immunomodulatory cytokines, including many of those assessed in the immunomodulatory panel used for cytokine analysis (*24, 46–58*). These pathways influence key immunomodulatory cytokines and chemokines such as TNFα, IL-1, IL-12p40, IL-10, IL-4, MCP-1 and MIP1β by activating transcription factors (*58*). To examine the role of these pathways in hMSCs yielding different therapeutic outcomes, the MAPK signaling pathway was screened in preliminary studies to identify differences in phospho-protein signaling between less therapeutic hMSCs (donor 1 and donor 2) and more therapeutic hMSCs (donor 3 and donor 4) in response to IL-1β exposure. To do so, we used a Luminex multiplexed immunoassay to quantify MAPK pathway signaling at 5-, 15-, and 60-minutes post IL-1β conditioning (Figure 5A, Sup. S4). Pathway signaling peaked at 15min (Sup. S4), revealing significantly increased phosphorylation in pJnk in the more therapeutic hMSCs compared to the less therapeutic hMSCs. However, there was no significant difference in the activating transcription factor (Atf)-2, heat shock protein (HSP)-27, p38, and c-Jun N-terminal kinase (JNK) in the more therapeutic hMSCs compared to less therapeutic hMSCs (Figure 5B). These results demonstrated that in more therapeutic hMSCs, MAPK signaling was elevated relative to less therapeutic hMSCs. More specifically, p-JNK phospho-protein levels in the MAPK pathway showed increased levels relative to less therapeutic hMSCs.

**Figure 5.**
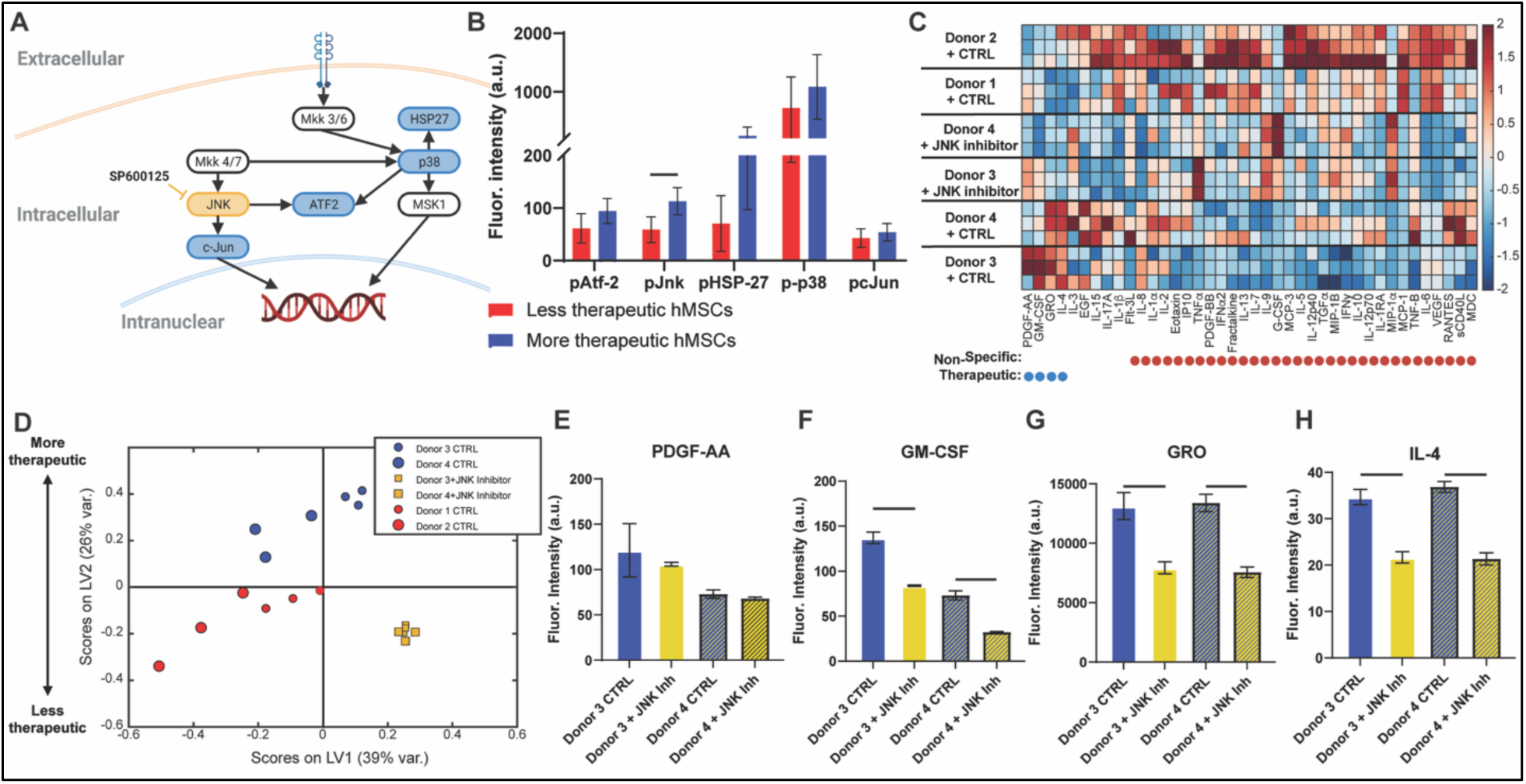
Paracrine signaling profiles of more therapeutic hMSCs treated with a p-JNK inhibitor (SP600125). (A) Signaling schematic of the MAPK pathway (ERK pathway is not included) in hMSCs with the inhibitor site (yellow) highlighted for the p-JNK phospho-protein signaling node. Quantification of MAPK phospho-protein signaling levels in all cell lines demonstrated increased overall phospho-protein signaling levels in the more therapeutic hMSCs relative to the less therapeutic hMSCs. (B) More therapeutic hMSCs demonstrated significantly increased expression of p-JNK phospho-protein signaling levels relative to less therapeutic hMSCs. (C) Quantification of immunomodulatory cytokines for more therapeutic hMSCs, with a p-JNK inhibitor applied, shows distinct shifts in overall cytokine signaling with addition of the inhibitor. (D) PLSDA analysis identified a profile of cytokines along LV2 that identified a distinct separation between more therapeutic hMSCs at top (blue) and more therapeutic hMSCs treated with inhibitors to the right (yellow). (E-H) Treatment with a p-JNK inhibitor yielded significantly less GM-CSF, GRO, and IL-4 secretion in more therapeutic hMSCs, relative to the more therapeutic hMSC control.

Because JNK phosphorylation was significantly elevated at 15 minutes post IL-1β conditioning, we next suppressed JNK signaling in more therapeutic hMSCs (donors 3 & 4) using the small molecule inhibitor SP600125 (p-JNK inhibitor) to determine the effect of JNK signaling on paracrine factor secretion. Treatment of more therapeutic hMSCs with the p-JNK inhibitor significantly reduced secretion of the key cytokines GM-CSF, GRO, and IL-4, (therapeutic associated cytokines) and a similar trend was observed for PDGF-AA but was not significant (Figure 5C – H). The JNK inhibitor treatment shifted the paracrine secretion profile of the more therapeutic cells toward a less therapeutic profile separating on LV 2 (Figure 5D). These data implicate JNK signaling as a critical differential regulatory pathway related to the therapeutic efficacy of hMSCs.

These studies identified key differences between less therapeutic and more therapeutic hMSCs in the MAPK signaling pathways in response to an OA simulated microenvironment. The findings are consistent with previous literature which have demonstrated the role that increased MAPK and Akt signaling in MSCs yield therapeutic outcomes in arthritic conditions (*46, 51, 59–61*). However, in these prior studies much of the focus of the role of MAPK and Akt signaling has centered around their role in MSC proliferation, differentiation, exosome production, apoptosis and senescence. However, these studies have yet to identify the role these signaling pathways play in mediating the secretion of immunomodulatory cytokines (*51, 60, 62*). The ability to modulate MSC paracrine signaling using these small molecule intervention strategies presents a promising approach to potentiate current MSC therapeutics used in the clinical space (*63*).

### Validation study: hMSCs secretion correlates to OA therapeutic outcomes

The initial training data studies revealed hMSC heterogeneity in key secreted cytokines in response to a simulated OA microenvironment and identified a strong correlation between an hMSC donors secreted cytokine profile and *in vivo* therapeutic outcomes. Different hMSC donors exhibited more or less therapeutic effects in the *in vivo* structural outcomes (Figure 2). Similarly, there was a distinct separation between the more and less therapeutic donors following *in vitro* secretome analysis (Figure 3). Data from these initial studies were used to train a PLSR feature learning model to identify predictive attributes of hMSCs therapeutic efficacy (Figure 1). To validate the predictive accuracy of this model, a prospective series of validation studies were performed. Four additional donor hMSCs (donor 5, donor 6, donor 7, donor 8) were acquired from RoosterBio, and the data from the new cells were run through the feature learning predictive model pipeline.

Structural outcome measurements from the *in vivo* training data studies were analyzed using principal component analysis. The results showed that structural outcome measurements were separated based on therapeutic efficacy with principal component 1 accounting for ∼60% (Sup S5) of the variance. Along the PC1 axis, saline control and sham control samples had the lowest and highest PC1 scores, respectively. More therapeutic hMSC donors (donor 3, donor 4) were closest to the sham samples and saw the most improvement in structural parameters in comparison to less therapeutic hMSC donors (donor 1, donor 2). A PLSR model was trained using cytokines as the independent variable and donor PC1 scores as the dependent variable.

The validation study showed similar variability in donor specific hMSC secretome following IL-1β stimulation compared to the training data study (Figure 6A). The secretome profile data from each additional donor hMSCs were projected onto the predictive PLSR model trained on the training data. The predicted composite disease scores were indicative of therapeutic efficacy (Model Projection & Predictions, Figure 1). The model projections prospectively revealed that the hMSCs from the new donors separated along LV axis 1 which was correlated to therapeutic efficacy (Figure 3B) thereby predicting less (donor 5, donor 6) and more therapeutic (donor 7, donor 8) donors (Figure 6B). Analysis of the secreted cytokine levels revealed significantly higher levels overall of all cytokines in the less therapeutic hMSCs compared with the more therapeutic hMSCs donors (Figure 6A). TNFα, MCP-1, G-CSF, and MIP-1 had significantly higher expression in the less therapeutic hMSCs in comparison to the more therapeutic hMSCs (Figures 6C-F).

**Figure 6.**
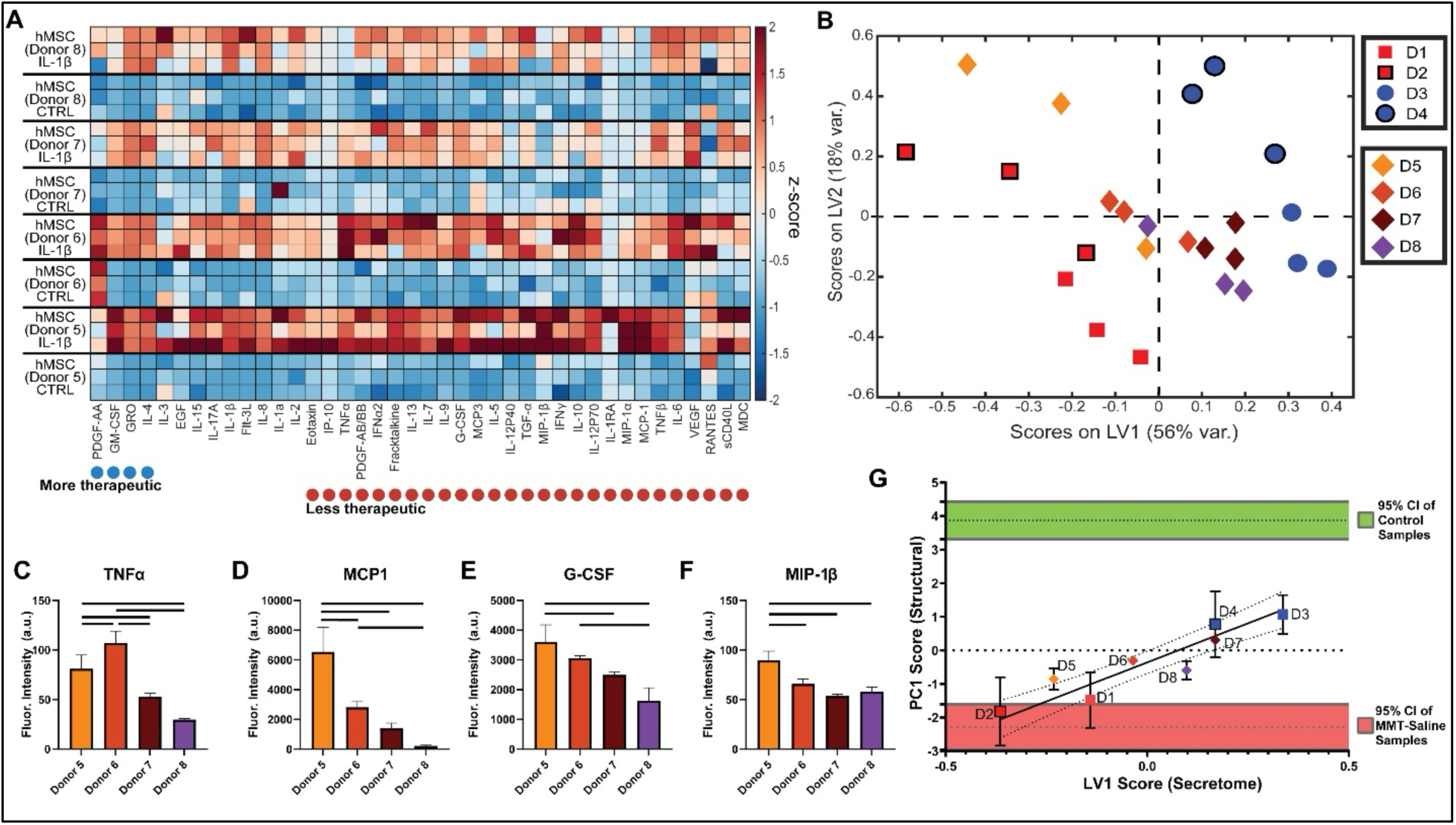
Validation study *in vitro* paracrine response of hMSCs in an OA simulated microenvironment. Multiplexed immunoassay analysis of 41 cytokines secreted from 4 new donor hMSCs in IL-1β conditioned media was completed for the validation studies. (A) Results showed increased expression of all cytokines in response to inflammatory stimulation in all hMSC donors. (B) New donors were projected on to the Initial PLSR model which identified a distinct separation between less therapeutic (red) and more therapeutic hMSCs (blue). Similarly, the new donors separated along LV1 with donor 5, donor 6 separating with the less therapeutic and donor 7 and donor 8 separating with the more therapeutic. Variability accounted for in each LV is included on respective axes labels. Donors 5 and 6 displayed higher levels of all cytokines, specifically (C) TNF⍺, (D) MCP1, (E) G-CSF, and (F) MIP-1β displayed significantly higher secretion levels relative to donors 7 and 8. (G) Displays linear relationship between PC1 structural scores and the LV1 secretome scores from the training data with donors 5, 6, 7, 8 secretome scores plotted to predict expected structural therapeutic efficacy.

Next, we conducted a linear regression analysis using the principal component 1 scores (PC1) for the training data (structural) along with the predicted PC1 scores for the validation data set against the LV1 scores (secretome) of all donors (Model projections and Prediction, Figure 1). The results demonstrated a positive linear relationship between the PC1 structural outcome scores and the LV1 secretome scores (Figure 6G). This framework showed the link between *in vitro* secretome profile and potential therapeutic efficacy for screening MSC donors in future *in vivo* validation studies.

### *In-Vivo* Therapeutic efficacy of hMSCs is predicted by *in-vitro* secretome response to IL-1β

Representative donor hMSCs were chosen from the less and more therapeutic groups for additional *in vivo* validation. Predicted less (donor 6) and more (donor 7) therapeutic hMSCs were administered intra-articularly in an *in vivo* OA preclinical study to prospectively validate the predictive capacity of the model. A z-scored heatmap of all of tibial structural parameters from microCT analysis shows increased expression of all outcomes in comparison to sham samples (Figures 7A). Exposed bone area and cartilage lesion volume was trending lower in the more therapeutic donor treated group with some samples not having lesion in comparison to the MMT-saline and less therapeutic donor group. However, there was variability across the samples and thus no statistical difference between the groups (Figure 7A).

**Figure 7:**
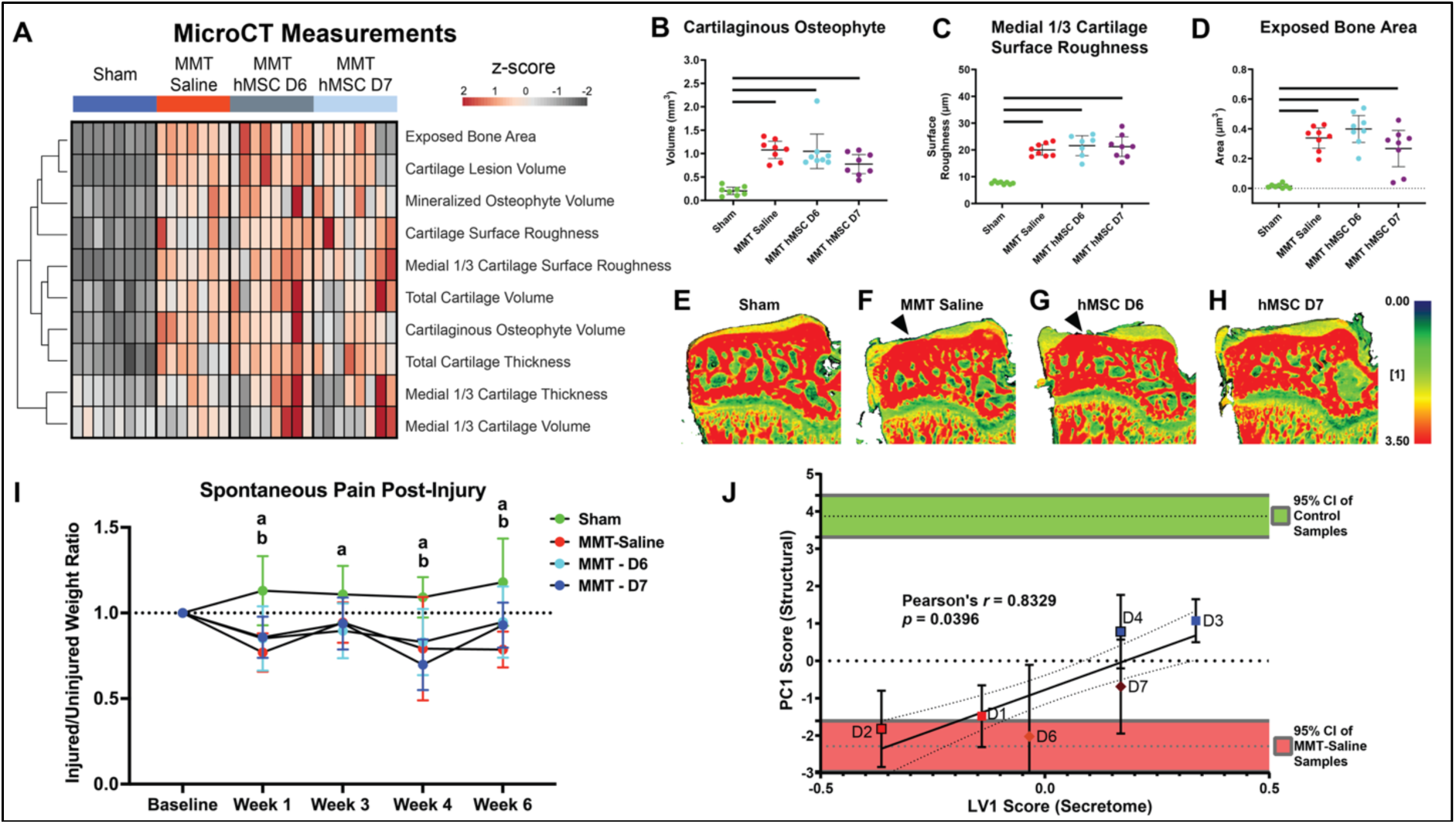
Validation study: Effect of predicted more and less therapeutic hMSCs on structural outcomes in the total and medial one-third (1/3) region of the medial articular cartilage in MMT joints. (A) Overall, all structural parameters increased in the MMT groups in comparison to sham. (B) Cartilaginous osteophyte volume, (C) medial 1/3 cartilage surface roughness, and (D) exposed bone significantly increased in all MMT groups relative to sham. Representative density heatmap images from contrast enhanced microCT revealed observable differences in structural outcomes. There were no structural cartilage lesions in the (E) sham group compared to the (F) MMT saline and (G) hMSC donor 6 groups (black arrowhead indicates lesion/exposed bone). (H) Donor 7 did not show lesion formation in some tibial plateaus, but these results were variable. (I) Spontaneous pain behaviors were significantly less in the sham group (represented by ‘a’) at all timepoints. (J) Displays correlation between PC1 structural scores and the LV1 secretome scores from the training and validation data. ‘b’ Represents significant differences (p < 0.05) between individual groups with each timepoint. ‘__’ Bars indicate significance (p < 0.05) between groups.

Cartilaginous osteophyte, mineralized osteophyte volume, and exposed bone area trended lower in the more therapeutic hMSCs but was not significant (Figure 7A - D). Similarly, representative cartilage thickness heatmap images from enhanced microCT revealed observable differences in structural outcomes (Figures 7E-H). There were no exposed bone lesions in the sham samples. The MMT-saline and less therapeutic hMSC (donor 6) treated group showed cartilage degradation and clear lesions in the articular cartilage extending to the subchondral bone (Figure 7E-G). Qualitatively in the more therapeutic hMSC (donor 7) treated group it appeared to demonstrate less cartilage degradation and lesions as displayed in the representative image (Figure 7H). But quantitatively there was no significant difference as a result of variability in the presence of lesions within the hMSC (donor 7) treated group. Spontaneous pain behavior was assessed longitudinally using dynamic weight bearing measurements. The injured to uninjured hindlimb weight ratio was significantly higher in the sham group compared to all MMT groups at all timepoints post-MMT surgery (Figure 7I). At 6 weeks post injury the injured to uninjured hindlimb weight ratio was significantly lower, indicating more less weight bearing on the injured limb, in the MMT-saline group compared to hMSC donor 6 treated group and trended lower compared to the hMSC donor 7 treated group (Figure 7I). In both hMSC treated groups the weight ratio returned to baseline levels at 6 weeks post injury in comparison to the MMT-saline group which was significantly lower indicting increased pain (Figure 7I). Injury triggered pain behaviors while the hMSC treatments exhibited protective effects in reducing pain behaviors 6 weeks post injury.

Following evaluation of structural outcome from hMSC donors *in vivo* in the validation studies (Sup Figure S6, S7) we performed a correlation analysis with the training study data (Figure 7J). The analysis revealed a significant correlation (Pearson *r* = 0.8329; *p* = 0.0396) between the PC1 structural scores and the LV1 secretome scores from our training and validation studies data (Model Validation, Figure 1). The more therapeutic donors had PC1 scores that trended closer to the 95% CI of the control samples while the less therapeutic donors had PC1 scores within the 95% CI of the MMT-saline samples. As the hMSC treatment was administered at 3 weeks post MMT, at a time when OA was already developed, these data suggest that the therapeutic donor MSCs are able to attenuate further progression of the disease but are not able to regenerate the tissue back to the naïve control state.

Together, these results demonstrate the high predictive capacity of our feature learning model to identify the therapeutic potential of individual donors hMSCs. Using this predictive feature learning model framework, we were able to classify hMSCs therapeutic potential based on their paracrine secretion of immunomodulatory cytokines and validate these predictions in a prospective *in vivo* validation study. Overall, the results revealed more protective effects in the predicted more therapeutic samples in comparison to the predicted less therapeutic samples. The results suggest variable therapeutic benefits of hMSCs can be prospectively predicted in our trained feature learning model pipeline and identifies a promising approach to optimize cell therapy for clinical translation and advance regenerative treatment strategies.

## CONCLUSION

Clinical trials evaluating the efficacy of MSC therapeutics for OA treatment have yielded highly variable therapeutic findings; this has motivated the need for standardization of screening metrics, and manufacturing protocols, to yield more consistent therapeutic outcomes following treatment delivery. Screening metrics for clinical utilization of MSCs is typically limited to adherence to tissue culture plastic, trilineage differentiation and expression of standard MSC phenotypic cell surface markers. This absence of better screening metrics that can identify critical quality attributes of MSCs and other cell therapies is likely a major contributor to the high variability that has been reported in clinical studies (*64–66*). To address this knowledge gap, we investigated and identified cellular attributes that relate to therapeutic outcomes of hMSCs in OA. Signature profiles for secreted cytokines, RNA transcripts, and intracellular signaling phospho-proteins were identified (due to their role in paracrine signaling) that relate to therapeutic outcomes in a pre-clinical model of OA. Pharmacological intervention strategies were also evaluated to mediate the production of the therapeutic cytokine. While further study is warranted to identify the efficacy of these intervention strategies *in vivo* the current studies present a promising therapeutic strategy for potentiating hMSC treatment of OA. These novel findings motivate promising research directions that with further study could be translated into the clinic to better screen and more effectively treat patients using cell based therapeutics.

In the current study, bone marrow derived hMSCs from four unique hMSC donors were analyzed to identify a profile of secreted cytokines, RNA transcripts, and signaling phospho-proteins of hMSCs that relate to therapeutic outcomes of hMSCs in OA. The therapeutic potential of all hMSCs was assessed in a pre-clinical rat model (MMT) of established OA (after onset of structural damage), which is a clinically relevant scenario as patients commonly seek treatment once OA is developed and symptoms are readily evident. A modeling analysis was performed to categorize each individual donors hMSCs as more therapeutic and less therapeutic. Analysis of cytokine secretion in response to an OA microenvironment showed a highly targeted paracrine secretome profile of the identified more therapeutic hMSCs with increased secretion of GM-CSF, GRO, IL-4, and PDGF-AA. Furthermore, RNA-Seq identified unique gene expression profiles of more therapeutic hMSCs, relative to less therapeutic hMSCs; pathway analysis showed significant differences in MAPK and Akt pathway signaling. Analysis of phospho-signaling in the MAPK pathway showed significant differences in p-JNK between more therapeutic and less therapeutic hMSCs. Data from these initial studies were used to train a feature learning model to predict therapeutic efficacy of additional hMSC donors which were then evaluated in a validation study (Figure 1). Comparably, in the validation study there was variability in hMSCs response to IL-1β stimulation. The feature learning model projections prospectively revealed hMSCs from the additional donors separated along LV1 which was predicted to be associated with therapeutic efficacy thereby predicting less and more therapeutic donors. Representative images from µCT qualitatively showed cartilage degradation and lesions in the MMT surgery groups compared to sham while the more therapeutic treatment group displayed less cartilage degradation. These results support the predicted structural outcomes from the validation study identifying the significant correlation and highlights the validity of the predictive power of the feature learning model.

The results supported our hypothesis that the feature learning model would predict therapeutic efficacy of the donors. The hMSC secretome were highly distinct between donor hMSCs which supports previous studies indicating the heterogeneity and variability of MSCs from different donors. Using our feature learning model pipeline, we were able to classify an hMSC donor’s therapeutic potential based on their paracrine secretion of immunomodulatory cytokines. Consistently identifying therapeutic donors will optimize regenerative cell therapy treatment for improved clinical translation. The findings indicate variable therapeutic benefits of hMSCs that can be prospectively predicted using a trained feature learning predictive model. hMSC cellular attributes were identified that relate to efficacious therapeutic outcomes; these provide critical quality attributes to prospectively identify therapeutic efficacy and identify targets to license or modulate the therapeutic potential of less therapeutic MSCs and ensure a highly consistent therapeutic product. Overall, this study increases the fundamental knowledge of hMSCs as therapeutics for use in OA, provide therapeutic targets, and a framework for identifying therapeutic hMSCs that are translatable into the clinic.

## MATERIALS AND METHODS

### Training Data and Validation Studies

#### hMSC culture and characterization

hMSCs derived from bone marrow were obtained from Emory Personalized Immunotherapy Core (EPIC) at Emory University and RoosterBio (MSC-004; RoosterBio, Inc; Frederick, MD, USA). EPIC hMSCs were cultured in Mesenchymal Stem Cell Basal Medium (PT-3238; Lonza; Basel, Switzerland) supplemented with 10% heat-inactivated FBS (S11110H; Atlanta Biologicals; Oakwood, GA, USA), 1 mM L-glutamine (SH3003401; HyClone; Logan, UT, USA), and 100 μg/mL Penicillin/Streptomycin (B21110; Atlanta Biologicals) and sub-cultured at 80% confluency. RoosterBio hMSCs were cultured in RoosterNourish-MSC (KT-001; RoosterBio, Inc) media supplemented with 2% RoosterBooster (SU-003; RoosterBio, Inc) and sub-cultured at 80% confluency. hMSC phenotypes for all donor hMSCs were confirmed by adipogenic, chondrogenic, and osteogenic differentiation. Flow cytometry was also used to characterize the hMSCs to confirm that all donor cells expressed characteristic MSC surface markers (CD73, CD90, CD105) and lacked hematopoietic markers (CD45, CD34, CD11b, CD79A, HLA-DR). Metadata was also collected for all hMSC donors (Table S1).

#### *In vivo* MMT model of OA

##### Training data studies

Animal care and experiments were conducted in accordance with the institutional guidelines of the VAMC and experimental procedures were approved by the Atlanta VAMC IACUC (Protocol: V004-15). Weight-matched wild type male Lewis rats (strain code: 004; Charles River), weighing 300-350 g, were acclimatized for 1 week after they were received. A surgical instability animal model, MMT, was used to induce OA, as previously described (*67*). hMSC therapeutics were assessed for their ability to prevent further development of established OA using a six week time course, with therapeutics injected at three weeks (corresponding to OA phenotype presentation) followed by animal takedown three weeks later at the six week end point (*46, 67*). All MMT animals received 50 μL intra-articular injections using a 25-gauge needle. Animals were injected with 1) HBSS (MMT/Saline; n=8), 2) 5×10^5^ hMSC/knee of donor 1 [MMT/hMSC(Donor 1); n=8], 3) 5×10^5^ hMSC/knee of donor 2 [MMT/hMSC(Donor 2), 4) 5×10^5^ hMSC/knee; n=8] of donor 3 [MMT/hMSC(Donor 3); n=7], or 5) 5×10^5^ hMSC/knee of donor 4 [MMT/hMSC(Donor 4); n=8]. The cell dose (5×10^5^ cells/knee) used for injection was matched with the dosage optimized in a prior study (*20*). Sham animals were not injected post-surgery (n=8). Animals were euthanized at 6-weeks post-surgery via CO2 asphyxiation. Left hindlimbs were dissected and fixed in 10% neutral buffered formalin. Muscle and connective tissues were removed from the hindlimbs. The femur was disarticulated from the tibia. Meniscus and residual soft tissue surrounding the medial tibial condyle were dissected and discarded.

##### Validation data studies

All procedures were conducted following IACUC protocol approval from the University of Oregon IACUC committee. Procedures were conducted as described above. Briefly, weight matched male Lewis rats (Charles River Laboratory) were randomly assigned to MMT or sham (n=8) surgery groups. Prior to injury, baseline spontaneous pain and limb function was assessed using Dynamic Weight Bearing testing (DWB 2.2.6, BIOSEB; validation study only). Rats then underwent either MMT or sham surgery (MCL transection only) according to group. Spontaneous pain and limb function analyses were performed again 1, 3, 4, and 6-weeks post-injury. Three weeks post-injury MMT rats received an intra-articular injection (50 μL saline or saline with 5×10^5^ cells suspension) according to randomly assigned treatment group. MMT-saline and two MMT-hMSC (Donor 6, Donor 7) treatment groups (n=8 per group). Three weeks post-injection, rats were euthanized and the hindlimbs collected for evaluation of PTOA structural pathology.

#### microCT quantitative analysis of articular joint parameters

##### Training data studies

Tibiae were immersed in 30% (diluted in PBS) hexabrix 320 contrast reagent (NDC 67684-5505-5; Guerbet) at 37 °C for 30 mins before being scanned (*68*). All samples were scanned using microCT (μCT 40, Scanco Medical) using the following parameters: 45 kVp, 177 μA, 200 ms integration time, isotropic 16 μm voxel size, and ∼27 min scan time (*68*). Native axial 2D tomograms were orthogonally transposed to sagittal and coronal views to streamline segmentation of local volumes of interest (VOIs) and subsequently evaluated to yield 3D reconstructions for all scanned samples. All microCT parameters (articular cartilage, osteophyte, and subchondral bone) were evaluated as previously described (*47, 69*). For cartilage parameters, thresholding of 110 - 435 mg hydroxyapatite per cubic cm (mg HA/cm^3^) was used to isolate the cartilage from the surrounding air and bone. Furthermore, for bone parameters, thresholds of 435 - 1200 mg HA/cm^3^ were implemented to isolate bone from the overlying cartilage. Samples were saved for histological processing following μCT imaging (S6A).

##### Validation data studies

All procedures were conducted as described above. Briefly, tibiae were immersed in 37.5% Conray (iothalamate meglumine) contrast reagent diluted in 62.5% ion-free PBS at 37°C for 30 mins before scanning. Tibiae samples were scanned in randomized order using a vivaCT80 (Scanco Medical) with the following parameters: 45 kVp, 177μA, 800 ms integration time, 16 μm voxel size, and ∼30min scan time. For evaluation, cartilage was isolated and analyzed the from surrounding air and bone. While bone was isolated and analyzed from the overlaying cartilage. Samples were saved for histological processing following μCT imaging (S6B).

#### MATLAB articular cartilage surface roughness analysis

Serial 2D sagittal images of the proximal tibiae were analyzed using a customized algorithm in MATLAB (MathWorks) to quantify surface roughness, lesion volume, and full-thickness lesion area (*48*). Images were processed to segment for the articular cartilage surface and the subchondral plate interface using intensity thresholding. The 3D articular cartilage surface rendering was fitted along an idealized 3D polynomial surface: fourth order along the ventral-dorsal axis and second order along the medial-lateral axis. The root mean square difference between the generated (actual) and polynomial fitted (predicted) surfaces was the measure of cartilage layer surface roughness. Lesion volume was calculated as the volume represented by the discrepancy in polynomial fit (predicted) surface and actual cartilage surface, anywhere that > 25% of total (predicted) cartilage thickness was exceeded. The exposed bone area was the sum of the area on the tibial condyle where no cartilage layer was present. Cartilage and bone segmentation proximity tolerance was 3 voxels (anything 3 voxels or fewer separation is considered to be no cartilage present). Surface roughness, lesion volume and full-thickness lesion area were calculated for full and medial 1/3 region of the articular cartilage.

#### *In vitro* OA simulated microenvironment

##### Training data studies

Matching donor hMSCs from the *in vivo* MMT model, were utilized *in vitro*. hMSCs were sub-cultured to 80% confluency in complete Lonza for hMSC donors donor 1 and donor 2 and Rooster medium for hMSC donors donor 3 and donor 4 in 12-well plates and cultured at 37 °C, 5% CO_2_. Each of the four hMSC donors were independently conditioned with 20 ng/mL IL-1β conditioned media (FHC05510; Promega; Madison, WI, USA; +IL-1β). IL-1β was used to model the OA inflammatory environment in the current study as it is a major pro-inflammatory modulator in OA (*24*). IL-1β concentrations were selected based on prior experiments and preliminary data (*49, 50*). The immunomodulatory cytokine content of all samples were characterized using a bead based multiplex immunoassay, Luminex Cytokine/Chemokine 41 Plex Immunomodulatory Kit (HCYTMAG-60K-PX41; EMD Millipore Corporation; Burlington, MA, USA). Median fluorescent intensity values were read out using Luminex xPONENT software V4.3 in the MAGPIX system. Conditioned media was added to hMSCs at day 0 followed by a 24-hr conditioning period in monolayer culture with media collection (Luminex cytokine analysis) and cell lysate collection (RNA-Seq analysis) at the study end point. Samples were stored at −80 °C until analysis was performed.

##### Validation data studies

Bone marrow derived hMSCs from four new donors were purchased from RoosterBio (Frederick, MD, USA) and cultured as described above. Briefly, the hMSCs were each sub-cultured for 24 hours in control media (K82016; RoosterNourishTM-MSC-XF media, RoosterBio) or conditioned media (stimulated with IL-1β 20 ng/ml). The media was collected and analyzed using a multiplexed immunoassay Cytokine/Chemokine 41 Plex Immunomodulatory kit (Luminex) to evaluate hMSCs secretome.

#### hMSC cytokine analysis

Cytokines were quantified for all donors with IL-1β conditioning (n=3). Loaded samples (2.03 μL) were determined to be within the linear range of detection of the MAGPIX (MAGPIX-XPON4.1-CEIVD; EMD Millipore Corporation) system. Cytokines were quantified using a bead based multiplex immunoassay, Luminex Cytokine/Chemokine 41 Plex Immunomodulatory Kit (HCYTMAG-60K-PX41; EMD Millipore Corporation). Median fluorescent intensity values were read out using Luminex xPONENT software V4.3 in the MAGPIX system. Background subtraction was performed using read out values 20 ng/mL IL-1β conditioned media. This assay was performed equally for all donors within the training data studies and the validation data studies.

#### hMSC RNAseq analysis (*Training Study only*)

RNA was isolated from hMSCs using the Qiagen RNeasy kit (217804; Qiagen; Hilden, Germany) according to the manufacturer’s protocol and submitted to the Georgia Tech Molecular Evolution Core for sequencing. Quality control was run on all samples using a bioanalyzer to determine that the RNA Integrity Number (RIN) was greater than 7 for all samples. A NEBNext Poly(A) mRNA Magnetic Isolation Module (E7490S; New England Biolab; Ipswich, MA, USA) and NEBNext Ultra II Directional RNA Library Prep Kit (E7760; New England Biolab) were used to generate libraries for sequencing. Sequencing was performed using the NovaSeq 6000 Sequencing System to obtain a sequencing depth of 30-40 million reads per sample. Samples were merged from four technical replicate lanes. Transcripts were aligned by Molecular Research LP (mrdnalab.com) using DNAstar Array Star and Qseq and reads were mapped to the homo sapiens (human) genome assembly GRCh38 (p14) from the genome reference database. For the read assignment, the threshold was set at 20bp and 80% of the bases matching within each read. Duplicated reads were eliminated and genes with less than 50 raw counts in at least 75% of the samples were removed from analysis. Non-coding genes were removed from the analysis. All gene counts were normalized using the DESeq2 R package (in R), available through Bioconductor. Gene ontology analysis on DEGs was conducted using PANTHER overrepresentation test with PANTHER 18.0 through the Gene Ontology resource (https://geneontology.org/) against all genes within the dataset. The Homo Sapiens Go Biological processes complete annotation set was used with Fisher’s exact test with FDR adjustment to compute significance of biological processes between More and Less therapeutic groups.

#### Gene set variation analysis (GSVA) *(Training Study Only)*

To establish differences in each gene set, we used GSVA to identify enrichment of gene sets across all donors (*45*). GSVA is an igene set enrichment method that detects subtle variations of pathway activity over a sample population in an unsupervised manner. The GSVA was conducted using the Molecular Signatures Database C2 and C7 gene sets (MSigDB) (*70*). Statistical differences in enrichment scores for each gene set between groups were computed by comparing the true differences in means against the differences computed by a random distribution obtained by permuting the gene labels 1000 times. False discovery rate (FDR) adjusted p-values were computed for detection of differences between donors with statistical significance set at FDR < 0.25 (*71*). GSVA was performed using the GSVA v1.36.1 in R (The R Foundation).

#### Partial least squares discriminant analysis (PLSDA) & Partial least squares regression analysis (PLSR)

PLSDA was performed in MATLAB (Mathworks) using a function written by Cleiton Nunes (Mathworks File Exchange) (*72*). This approach accounts for the multivariate nature of the data without overfitting (*73, 74*). Prior to inputting the data into the algorithm, all data was z-scored. Secreted cytokines level read outs (from the pharmacological intervention studies) were used as the independent variables and the different hMSC donors/treatments were used as the binary outcome variables. LVs in a multidimensional space (dimensionality varied by number of independent input variables) were defined, and the two primary LVs were used for orthogonal rotation to best separate groups in the new plane defined by LV1 and LV2. PLSR was performed using the same function as PLSDA but specifying for PLSR modeling. Secretome data for each donor was first z-scored prior to the analysis. Z-scored secretome comprised of individual cytokine measurements were used as the independent variables and the average of the first principal component scores from the *in-vivo* study was used as the dependent response variable. This model was then used for predicting therapeutic efficacy of new hMSCs *in-vivo*. Within the model projections stage the function “plspred” by Cleiton Nunes was used with new z-scored hMSC secretome data as the new samples for prediction and the prior PLSR model as the calibration parameter.

#### Principal component analysis (PCA) of in-vivo microCT measurements

PCA was performed in MATLAB (Mathworks) using the built-in *pca* function. *In-vivo* microCT measurements for each group were first normalized to the average of the saline control group for each respective measurement and then z-scored. Final composite values (scores) were recorded for each group and later used as the dependent variable in creating a partial least squares regression model for in-vitro hMSC response to IL-1β.

#### Statistical analysis

All data is presented as mean ± SD. Weight matched samples were randomly assigned to experimental groups and each animal assigned a sample identification number. Investigators were blinded to the treatment group classifications throughout data collection and analysis. Significance for all microCT parameters was determined with one-way ANOVA with post hoc Tukey honest test for articular cartilage and subchondral bone parameters. Bonferroni correction was used for post hoc analysis for the exposed bone and osteophyte parameters due to their nonparametric nature. To determine significant differences between different hMSC conditions (donor vs. donor, +CTRL vs. +drug, more therapeutic vs. less therapeutic) for individual cytokines and phospho-proteins, two tailed t-tests were used with Bonferroni correction to account for the independent analysis of multiple groups. To quantify the relationship between RNA gene expression pathways and articular cartilage microCT outcomes, least squares linear regression models were generated and Pearson’s correlation coefficient, R, together with the p-value calculated from an F test (null hypothesis that the overall slope is zero) were reported. To assess the relationship between secretome profiles and structural outcomes, a correlational analysis was conducted. For all analysis, p < .05 was considered statistically significant. All analyses were performed using GraphPad Prism 10 (GraphPad Software; La Jolla, CA, USA).

## Acknowledgments

We thank Benjamin T. Tignor for his contribution to the completion of this study.

## Author contributions

Conceptualization: SMS, YCP, JMM, SB, LBW, NJW

Methodology: SMS, YCP, JMM, SB, NMP, AL, KL

Investigation: SMS, YCP, JMM, SB, NMP, AL, KL

Visualization: SMS, YCP, JMM, SB, NMP, AL, KL

Funding acquisition: LBW, NJW

Project administration: LBW, NJW

Supervision: LBW, NJW

Writing – original draft: SMS, YCP, JMM, SB, LBW, NJW

Writing – review & editing: SMS, YCP, SB, LBW, NJW

## Data and materials availability

All data are available in the main text or the supplementary materials.

