## Supplemental Tables and Figures for "A Feature Learning Model Identifies Predictive Attributes of Mesenchymal Stromal Cell Efficacy"

**Supplemental Figures**

| Donor # | Source | Sex | Age | Tissue | Passage | Donor Identification | Study |
|---|---|---|---|---|---|---|---|
| **1** | EPIC | F | 5 | Bone Marrow | 4 | Donor 1 | Training |
| **2** | EPIC | M | 4 – 20 | Bone Marrow | 4 | Donor 2 | Training |
| **3** | RoosterBio | M | 18 – 30 | Bone Marrow | 4 | Donor 3 | Training |
| **4** | RoosterBio | F | 26 | Bone Marrow | 4 | Donor 4 | Training |
| **5** | RoosterBio | M | 23 | Bone Marrow | 4 | Donor 5 | Validation |
| **6** | RoosterBio | F | 26 | Bone Marrow | 4 | Donor 6 | Validation |
| **7** | RoosterBio | M | 29 | Bone Marrow | 4 | Donor 7 | Validation |
| **8** | RoosterBio | M | 25 | Bone Marrow | 4 | Donor 8 | Validation |

**Table S1: Metadata for hMSC donors.** hMSC samples used in the experiments for this study

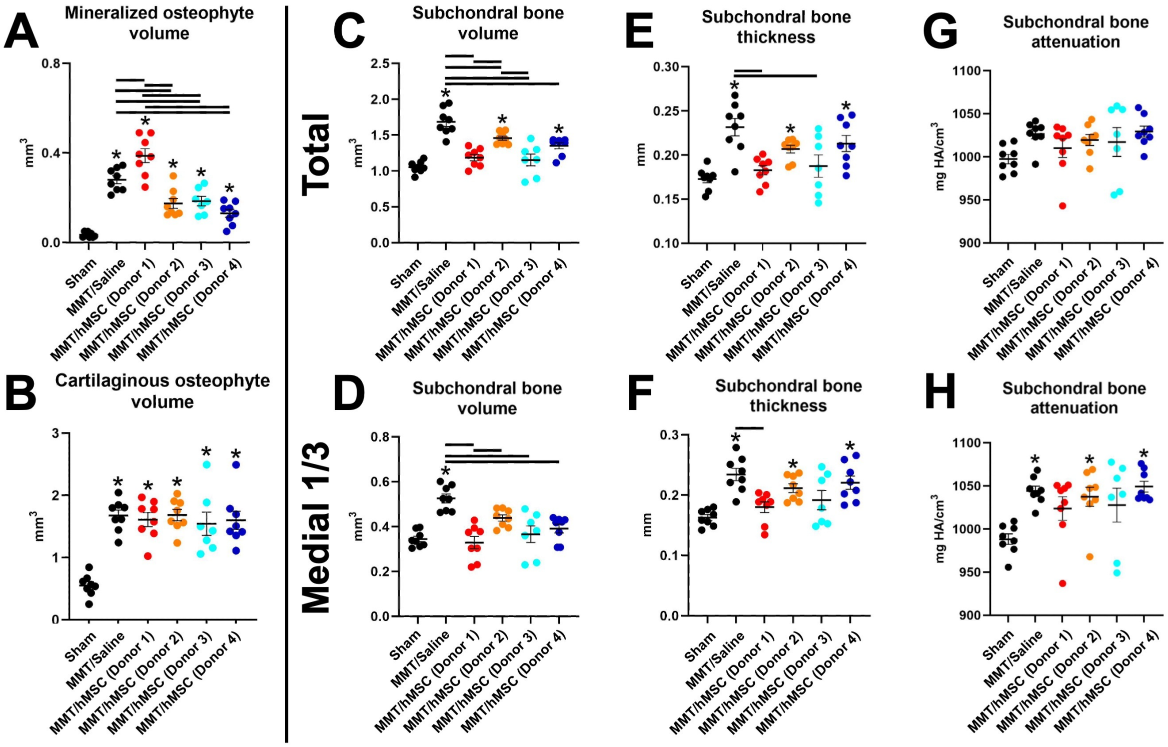


**Figure S1. Effect of hMSC donor heterogeneity on osteophyte and subchondral bone therapeutic outcomes in the total and medial ⅓ of the medial articular tibial cartilage in 6-week MMT joints.** (A) Represents mineralized osteophyte volumes which were significantly increased in all MMT groups relative to sham. (B) Cartilaginous osteophyte volumes revealed greater volumes in all MMT groups compared to sham, but no differences were noted between MMT groups. (C) Total subchondral bone volume showed MMT/Saline, MMT/hMSC (Donor 2), and MMT/hMSC (Donor 4) groups had significantly higher volume in comparison to sham. Quantitative analysis results from (D) subchondral bone volume in the medial 1/3 region, (E) total subchondral bone thickness (F) medial 1/3 region subchondral bone thickness (G) subchondral bone attenuation of the total tibial plateau and (H) subchondral bone attenuation in the medial 1/3 region. Data presented as mean +/- SD. n=7 for MMT/hMSC (Donor 3) and n=8 for all other groups. Asterisk (*) Represents significant differences ($p < 0.05$) between individual MMT groups and sham. Horizontal black bars indicate significance ($p < 0.05$) between individual MMT groups.

### Table S2: Excel File attached separately

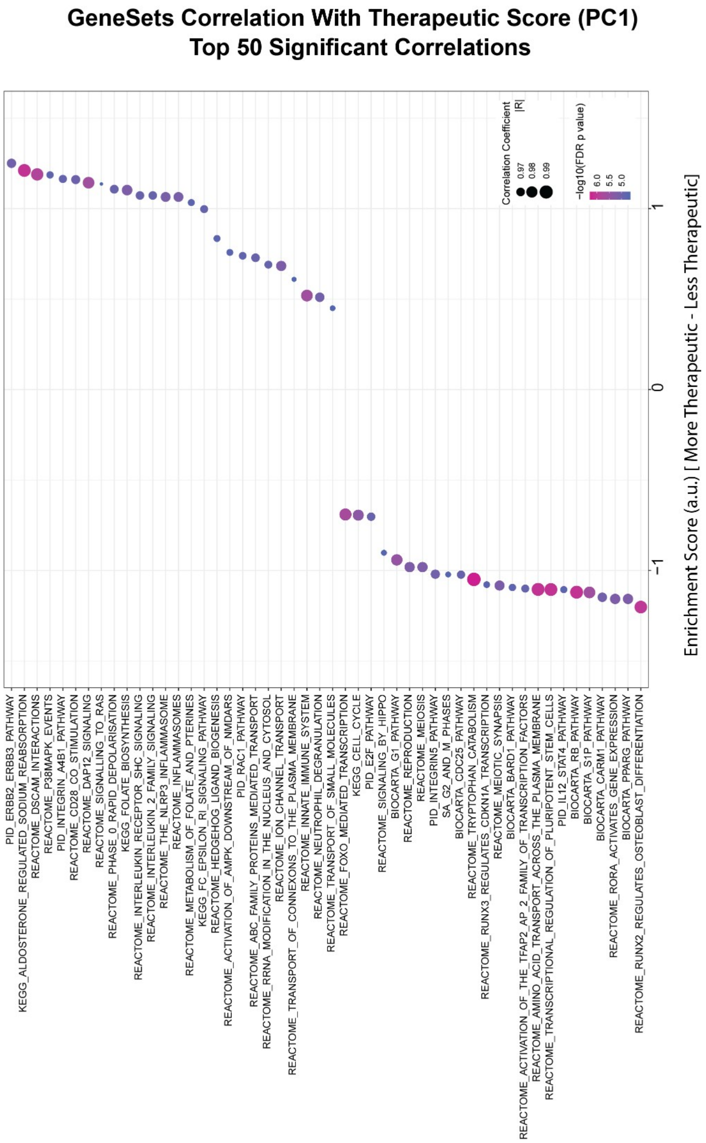


**Figure S2.Top 50 significant Pearson's correlation coefficient of therapeutic score (PC1) versus gene sets enrichment score identified by GSVA (rows) in hMSC.** The size of dots represents absolute value of the correlation coefficient (|R|) and the ones with negative correlation are noted with red asterisk (*).The color bar represents p-values (FDR adjusted) associated with the correlation slope.

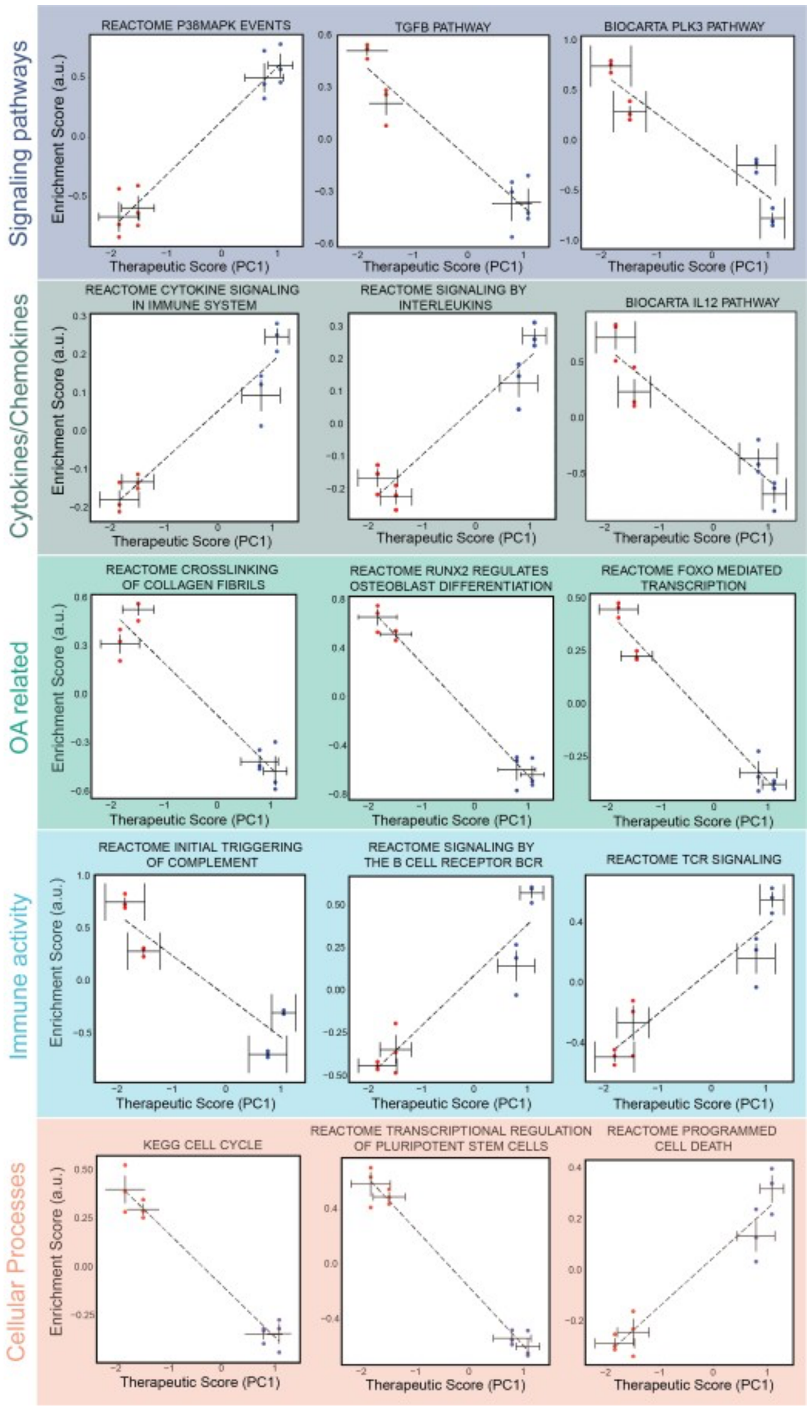


**Figure S3. Transcriptional signatures associated with signaling pathways, cytokines, OA related, immune activity, and cellular processes gene sets correlates with therapeutic scores.** Pearson's correlation analysis of therapeutic score (PC1) versus gene sets enrichment score identified by GSVA in hMSC (mean±SEM, N=3).

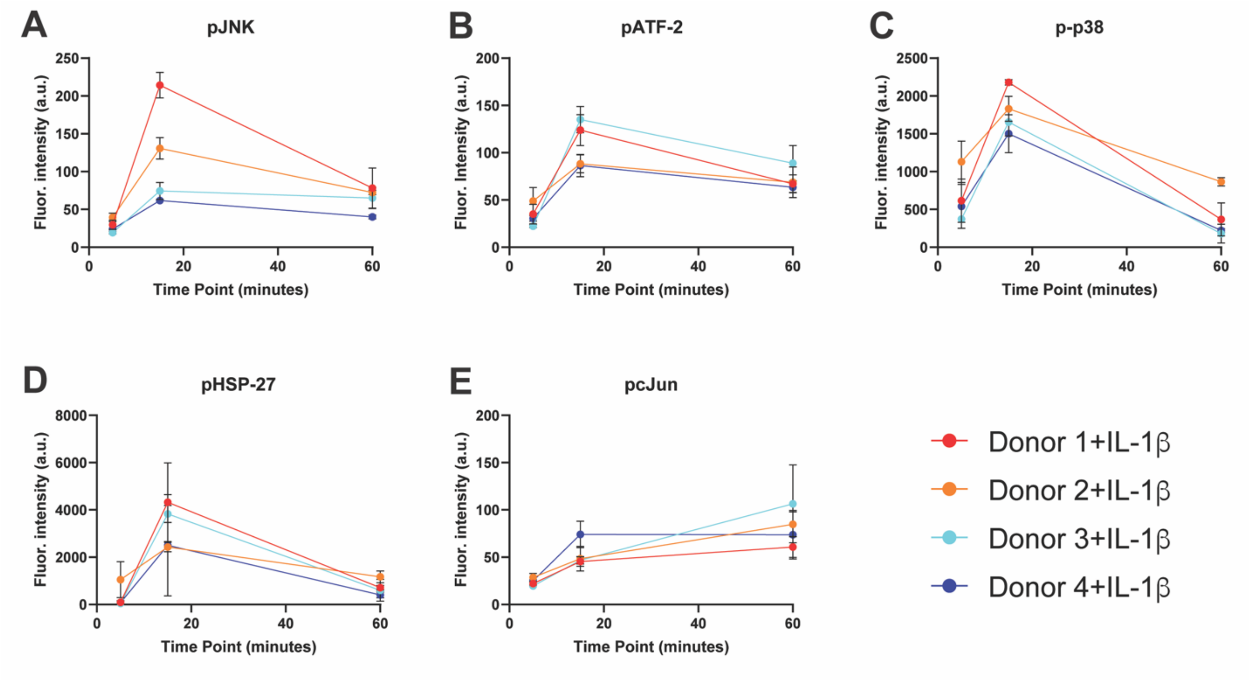


**Figure S4. Temporal analysis of target phospho-protein signals in the MAPK and Akt signaling pathways.** Quantitative analysis of (A) p-JNK, (B) p-Atf-2, (C) p-p38, (D) pHSP-27, and (E) pcJun phospho-protein signaling in the MAPK pathway revealed signaling spikes in 15 mins, relative to the 5- and 60-min time points.

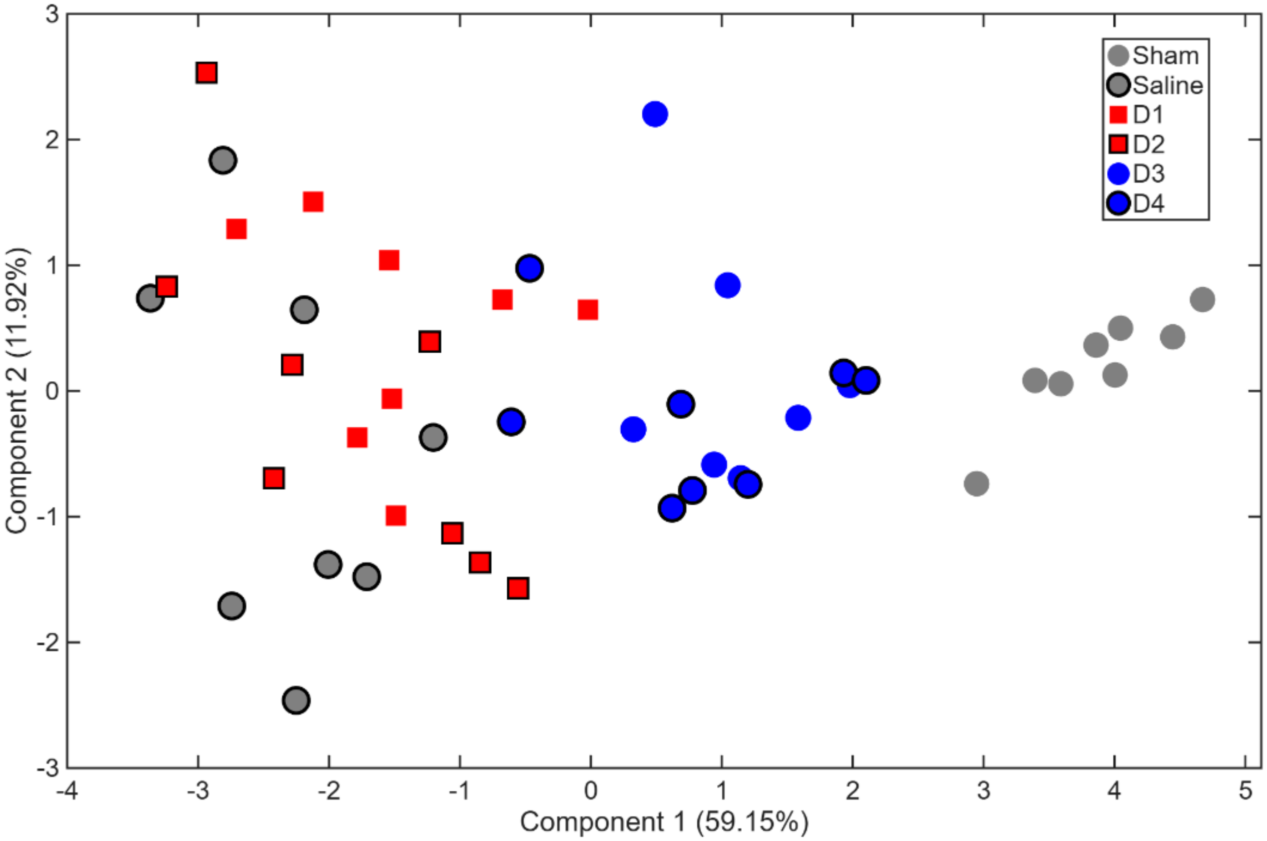


**Figure S5. Scatter plot of principal component 1 and 2 for *in-vivo* training study structural parameters.** Structural outcome measurements were separated through principal component analysis with principal component 1 accounting for 59.12% and principal component 2 accounting for 11.92%.

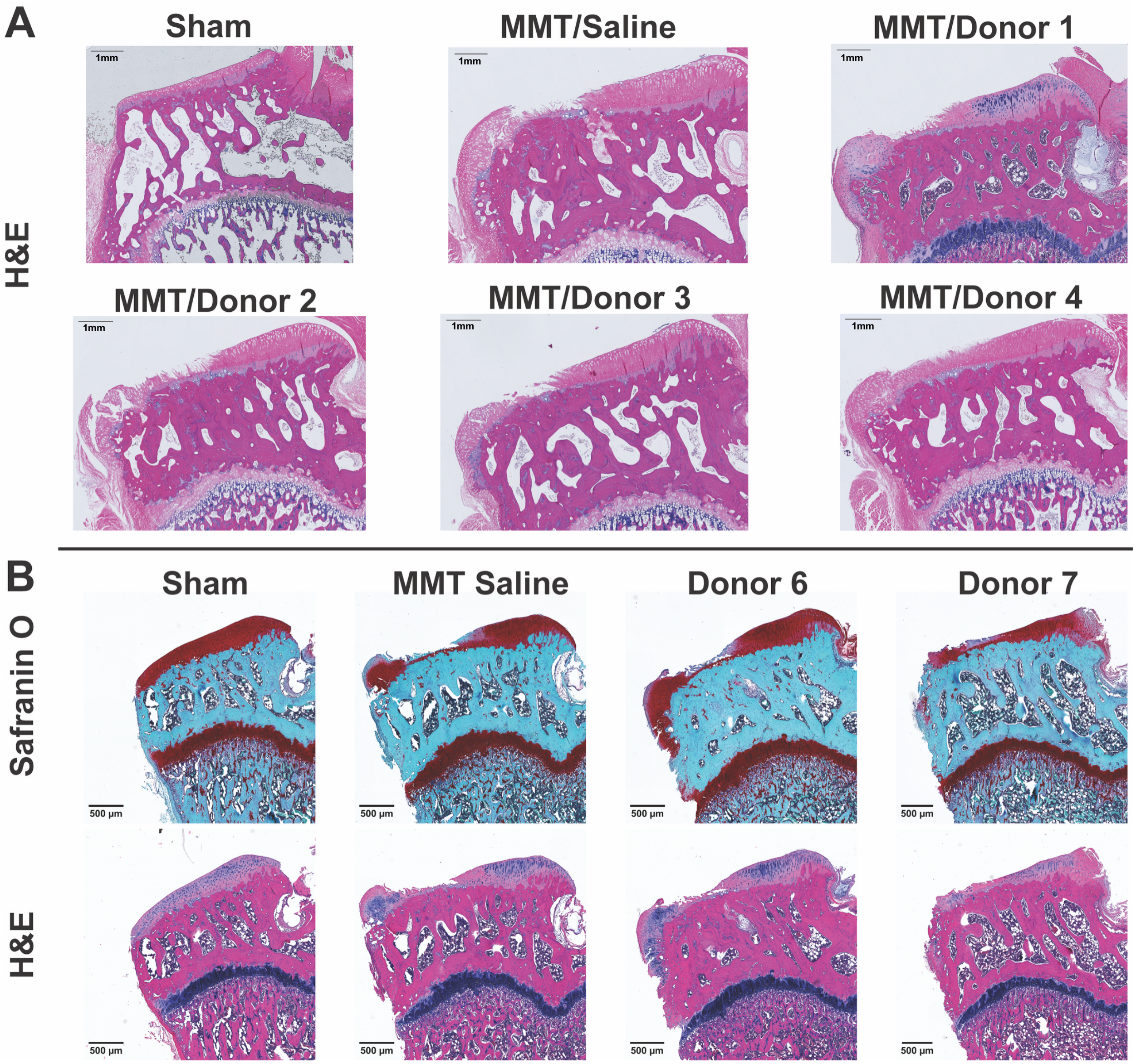


**Figure S6. Representative histology of training and validation study animals.** (A) Representative hematoxylin and eosin-y (H&E) staining of training study animals. (B) Safranin O and H&E staining of validation study animals following µCT imaging.

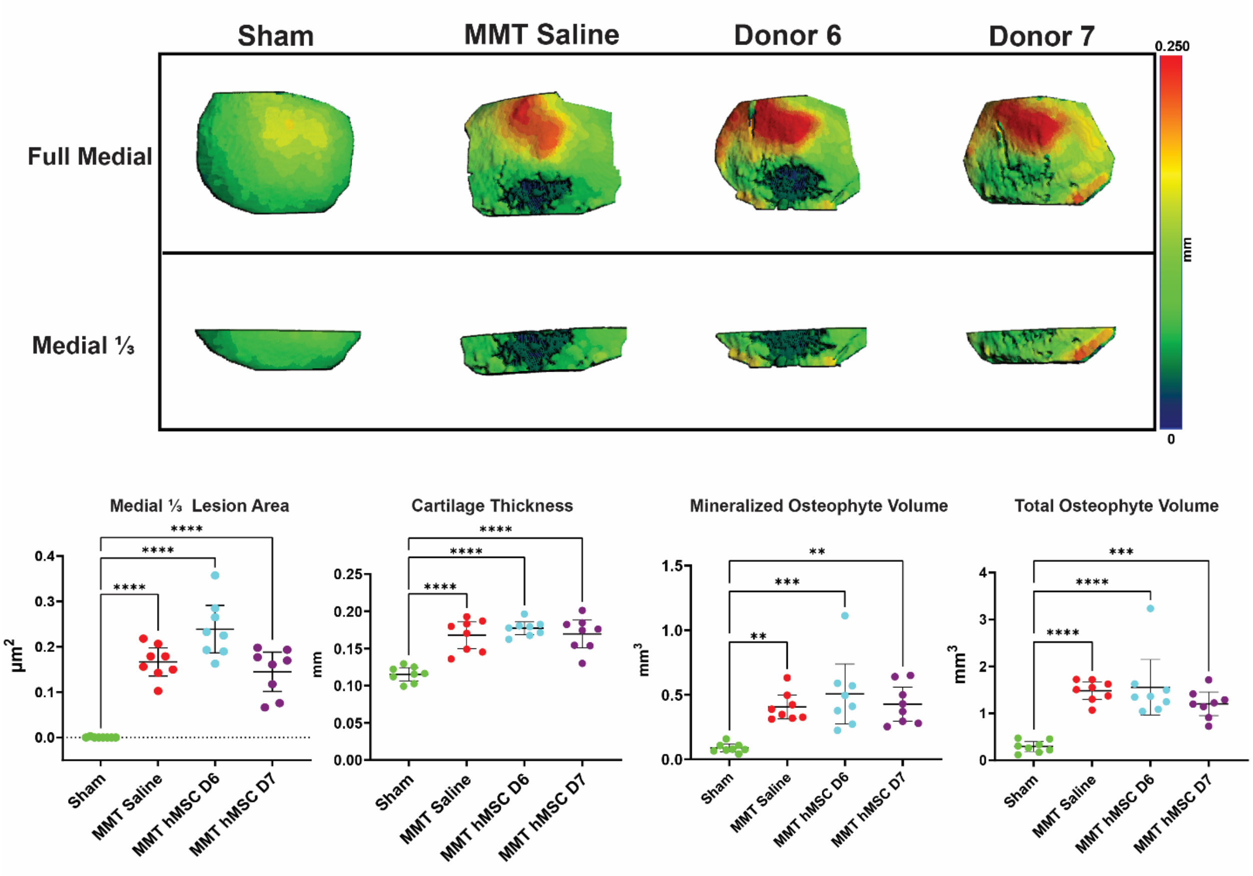


**Figure S7. Representative contrast enhanced μCT images and structural outcomes from the validation study.** Images shown display the full and medial 1/3 compartment of animals and effect of hMSC treatment on structural parameters.
